# Extracellular vesicle-associated cholesterol dictates the regenerative functions of macrophages in the brain

**DOI:** 10.1101/2022.12.23.521775

**Authors:** Sam Vanherle, Jeroen Guns, Melanie Loix, Fleur Mingneau, Tess Dierckx, Tim Vangansewinkel, Esther Wolfs, Paula Pincela Lins, Annelies Bronckaers, Ivo Lambrichts, Jonas Dehairs, Johannes V. Swinnen, Sanne G.S. Verberk, Mansour Haidar, Jerome J.A. Hendriks, Jeroen F.J. Bogie

**Author notes:** These authors contributed equally (Jerome J.A. Hendriks, Jeroen F.J. Bogie).

## Abstract

Macrophages play major roles in the pathophysiology of various neurological disorders, being involved in seemingly opposing processes such as lesion progression and resolution. Yet, the molecular mechanisms that drive their harmful and benign effector functions remain poorly understood. Here, we demonstrate that extracellular vesicles (EVs) secreted by repair-associated macrophages (RAMs) enhance remyelination *ex vivo* and *in vivo* by promoting the differentiation of oligodendrocyte precursor cells (OPCs). Guided by lipidomic analysis and applying cholesterol depletion and enrichment strategies, we find that EVs released by RAMs show markedly elevated cholesterol levels and that cholestserol abundance controls their reparative impact on OPC maturation and remyelination. Mechanistically, EV-associated cholesterol was found to promote OPC differentiation through direct membrane fusion. Collectively, our findings highlight that EVs are essential for cholesterol trafficking in the brain and that changes in cholesterol abundance dictate the reparative impact of EVs released by macrophages in the brain, potentially having broad implications for therapeutic strategies aimed at promoting repair in neurodegenerative disorders.

## Introduction

Chronic demyelinating diseases of the central nervous system (CNS) are characterized by loss of the protective myelin sheath, leaving axons and even complete neurons vulnerable to degeneration (Franklin and Ffrench-Constant, 2017). In early disease stages, endogenous repair processes are apparent, evidenced by lesional migration of oligodendrocyte progenitor cells (OPCs) and their successive differentiation into mature myelinating oligodendrocytes (OLNs) (Di Bello et al., 1999; Fancy et al., 2010; Franklin and Ffrench-Constant, 2017). However, as disease progresses, endogenous repair mechanisms frequently fail, resulting in the formation of chronically demyelinated lesions and widespread axonal and neuronal degeneration (Cantuti-Castelvetri et al., 2018; Hagemeier et al., 2012; Hanafy and Sloane, 2011; Wolswijk, 2002). These cytodegenerative events are anticipated to drive clinical deterioration in patients, often leading to irreversible physical, neurological, and cognitive incapacitation. Emerging evidence indicates that failure of remyelination relies on microenvironmental factors to which OPCs are exposed to in demyelinating lesions (Neumann et al., 2019; Wang et al., 2020; Woodruff et al., 2004). To date, however, the cell types, soluble factors, and molecular mechanisms involved remain poorly understood.

Dysfunction or imbalances in the macrophage phenotype are increasingly being acknowledged to drive demyelinating events and underlie failure of remyelination in the CNS (Bogie et al., 2020; Cantuti-Castelvetri et al., 2018; Marschallinger et al., 2020). In support of this notion, CNS lesions are characterized by abundant accumulation of disease-associated macrophages (DAMs), which are considered to promote demyelination and hamper remyelination through the release of inflammatory and toxic mediators that negatively impact OPCs, OLNs, and neuronal homeostasis (Bogie et al., 2020; Cantuti-Castelvetri et al., 2018; Marschallinger et al., 2020; McMahon et al., 2002; Nikić et al., 2011; Trapp et al., 1998). Whereas macrophages can acquire disease-resolving features, including the release of neurotrophic factors and clearance of inhibitory myelin debris (Bogie et al., 2020; Miron et al., 2013; Ruckh et al., 2012), the induction of such repair-associated macrophages (RAMs) appears to be transient. Accordingly, we and others demonstrated that, in time, perturbed lipid metabolism quenches and promotes the formation of RAMs and DAMs, respectively (Bogie et al., 2020; Cantuti-Castelvetri et al., 2018). Despite being closely associated with lesion progression and resolution, the harmful and benign effector mechanisms of RAMs and DAMs are ill-defined. Identifying factors that underlie the effector functions of RAMs and DAMs is essential for our understanding of lesion progression in chronic demyelinating disorders and the development of pro-regenerative therapies.

Extracellular vesicles (EVs) are nanosized particles surrounded by a lipid bilayer membrane that transport biologically active molecules such as proteins, nucleic acids, and lipids from donor cells to distant and neighboring recipient cells, thereby affecting recipient cell physiology (Kalluri and LeBleu, 2020; Vanherle et al., 2020). Emerging evidence indicates that EVs are associated with degenerative and regenerative events in the CNS (Guo et al., 2020; Herman et al., 2022; Hervera et al., 2018), and that modified EVs represent a promising therapeutic tool to promote CNS repair (Cooper et al., 2014; Wang et al., 2019; Zhang et al., 2021). Here, we report that EVs released by RAMs enhance OPC maturation *in vitro* and myelin regeneration in *ex vivo* and *in vivo* models that mimic remyelination. Guided by lipidomic analysis, we find that the pro-regenerative impact of EVs released by RAMs is ascribed to increased cholesterol abundance in these EVs. Accordingly, enrichment of cholesterol in EVs released by DAMs markedly enhances their regenerative properties. Finally, we provide evidence that direct membrane incorporation of EV-associated cholesterol via membrane fusion underlies the regenerative impact of EVs released by RAMs on OPC maturation and remyelination. Collectively, our findings highlight that EVs are essential for cholesterol trafficking and that changes in cholesterol abundance dictate the benign impact of EVs released by macrophages in the CNS.

## Materials and methods

### Antibodies and chemical reagents

The following antibodies were used for immunofluorescent/immunohistochemical stainings: anti-Flotillin 1 (1:1000, cat. #D2V7J, Cell Signalling Technology), anti-CD81 (1:1000, cat. #D502Q, Cell Signalling Technology), anti-Annexin A2 (1:1000, cat. #D1162, Cell Signalling Technology), anti-GRP94 (1:1000, cat. #D6X2Q, Cell Signalling Technology), anti-MBP (1:250 (*in vivo, ex vivo*) or 1:500 (*in vitro*), cat. #MAB386, Millipore), anti-O4 (1:1000, cat. #MAB1326, R&D systems), anti-Olig2 (1:100, cat. #AF2418, R&D systems), anti-CC1 (1:100, cat. #ab16794, Sigma-Aldrich), anti-NF (1:1000, cat. #ab8135, Sigma-Aldrich). Appropriate secondary antibodies were purchased from Invitrogen. Methyl-β-cyclodextrin (MβCD; 1, 2.5, or 5 mg/mL, cat. #C4555, Sigma-Aldrich) was used to deplete EVs from cholesterol or was complexed with cholesterol (7.3 mM, cat. #C3045, Sigma-Aldrich) to enrich EVs with cholesterol. Cytochalasin D (10 µM, cat. #C8273, Sigma-Aldrich), chlorpromazine hydrochloride (50 µM, cat. #C8138, Sigma-Aldrich), nystatin (25 µM, cat. #N4014, Sigma-Aldrich), imipramine hydrochloride (5 µM, cat. #I7379, Sigma-Aldrich), wortmannin (500 nM, cat. #W1628, Sigma-Aldrich), omeprazole (125, 250, 500 µM, cat. #O104, Sigma-Aldrich) and MβCD (2 mg/mL) were used to inhibit specific EV uptake pathways. T0901317 (10 µM, cat. #71810, Cayman Chemicals)

### Animals

Wild-type C57BL/6J mice were purchased from Envigo and were housed in a 12 h light/dark cycle with *ad libitum* access to water and a standard chow diet or specific formulations as indicated. All animal procedures were conducted in accordance with the institutional guidelines and approved by the Ethical Committee for Animal Experiments of Hasselt University (protocol numbers 201802, 201934K, 201967).

### EV production and isolation

Bone marrow-derived macrophages (BMDMs) were obtained as described previously (Bogie et al., 2017). Briefly, femoral and tibial bone marrow was isolated from 11-week-old female wild-type C57BL/6J mice and cultured in 14.5 cm petri dishes (Greiner, ref. 639161) at a concentration of 10 × 10^6^ cells/petri dish in RPMI1640 medium (Lonza) supplemented with 10% fetal calf serum (FCS; Biowest), 50 U/mL penicillin (Invitrogen), 50 U/mL streptomycin (Invitrogen), and 15% L929-conditioned medium (LCM) for 7 days at 37°C and 5% CO_2_. Next, cells were cultured at a density of 0.5 × 10^6^ cells/mL in RPMI1640 supplemented with 10% FCS, 50 U/mL penicillin, 50 U/mL streptomycin, and 5% LCM at a total of 13.5 × 10^6^ cells in T175 culture flasks (Greiner). Subsequently, BMDMs were treated for 6 h with vehicle, LPS (100 ng/µL, Sigma-Aldrich), or IL-4 (20 ng/µL, Peprotech) to induce naive macrophages (M0), DAMs, or RAMs, respectively, after which BMDMs were cultured in complete medium. The next morning, conditioned medium was collected, shortly centrifuged to remove detached cells and stored at -80°C. Next, BMDMs were cultured in EV-depleted culture medium (RPMI1640 supplemented with 10% FCS, 50 U/mL penicillin, 50 U/mL streptomycin, and 5% LCM, centrifuged for 16 h at 115,000 g, followed by 0.2 µm filtration) and medium was collected after 1 h (repeated four times). Thereafter, EVs were pelleted using a differential centrifugation method. Briefly, collected supernatant was consecutively centrifuged at 300 g for 10 min, 2,000 g for 10 min, and 10,000 g for 30 min, all at 4°C. Subsequently, supernatant was centrifuged for 3 h at 115,000 g and 4°C using the XPN-80 ultracentrifuge equipped with a Ti70 rotor (Beckman). EV pellets were resolved in 1 mL ice-cold PBS (Gibco) and further purified by means of size exclusion chromatography (SEC) using a chromatography column (Bio-Rad Laboratories) filled with 10 mL Sepharose CL-2B beads (GE Healthcare). Sequential fraction 3.5 until 5.5 mL was collected from the flow-through and further concentrated using the Amicon Ultra 0.5 mL 10 kDa filter (Millipore). To isolate EV-associated lipids, EVs were destroyed by hypo-osmotic shock and pelleted as previously described (Gabrielli et al., 2015). Lipids were extracted using chloroform and methanol (2:1 v/v), and subsequently, the lipid fraction was evaporated and dried for 1 h at 50°C. Next, EV-associated lipids were resuspended in PBS at 40°C and sonicated to obtain small vesicles.

### Nanoparticle tracking analysis

To determine the size and concentration of BMDM-derived EVs, Nanoparticle Tracking Analysis (NTA) was performed using the NanoSight NS300 system (Malvern Panalytical), equipped with a 532 nm laser. Samples were diluted in PBS over a concentration range between 20 and 50 particles per frame. All settings were manually set at the start of the measurements and maintained during all acquisitions: camera level 14, camera gain 1, pump rate 80, viscosity 1. A minimum of three recordings of 30 s was recorded and analyzed using the NTA software 3.0 with default settings.

### Western Blot

EVs were lysed in ice-cold Radio-Immunoprecipiration Assay (RIPA) buffer (150 mM NaCl, 50 mM Tris, 1% SDS, 1% Triton X-100, and 0.5% sodium deoxycholate) supplemented with protease-phosphatase inhibitor cocktail (Roche). Samples were separated by electrophoresis on a 10% SDS-PAGE gel and were transferred onto a PVDF membrane. Blots were blocked using 5% dried skimmed milk powder (Marvel) in 1x PBS-0.1% Tween-20 (PBS-T), incubated overnight with the relevant primary antibody, followed by incubation with the appropriate HRP-conjugated secondary antibody. Chemiluminescent signals were detected with the Amersham Imager 680 (GE Healthcare Life Sciences), using the enhanced chemiluminescence (ECL) Plus detection kit (Thermo Fisher Scientific).

### Cuprizone-induced acute demyelination *in vivo* model

To induce acute demyelination, 9-week-old male wild-type C57BL/6J mice were fed a diet containing 0.3% (w/w) cuprizone (bis[cyclohexanone]oxaldihydrazone, Sigma-Aldrich) mixed in powdered standard chow *ad libitum* for 5 weeks. Subsequently, mice were stereotactically injected intracerebroventricular with vehicle or BMDM-derived EVs. Briefly, mice were anesthetized by intraperitoneal injection with a mixture of 10% w/v ketamine (Anesketin, Dechra), 2% w/v xylazine (Rompun, Bayer), and PBS (1 mg ketamine and 0.12 mg xylazine per 10 g body weight, dose volume 0.1 mL/10 g). Next, vehicle or BMDM-derived EVs (1 × 10^9^ EVs) were unilaterally (left hemisphere) injected (AP 1.1 mm, ML -0.7 mm, DV -2.0 mm, relative to bregma) using a 10 µL Hamilton syringe at a speed of 1 µL/min. After injection, the needle was kept in place for an additional 2 min. After injection, cuprizone diet was withdrawn to allow spontaneous remyelination. 5 or 6 weeks after the start of cuprizone diet, mice were sacrificed, and tissue was collected for histological and biochemical analysis.

### Cerebellar brain slice cultures

Cerebellar brain slices were obtained from wild-type C57BL/6J mouse pups at postnatal day 10 (P10), as described previously (Hussain et al., 2011; Meffre et al., 2015). Brain slices were cultured in MEM medium (Thermo Fisher Scientific) supplemented with 25% horse serum (Thermo Fisher Scientific), 25% Hank’s balanced salt solution (Sigma-Aldrich), 50 U/mL penicillin, 50 U/mL streptomycin, 1% Glutamax (Thermo Fisher Scientific), 12.5 mM HEPES (Thermo Fisher Scientific), and 1.45 g/L glucose (Sigma-Aldrich). To induce demyelination, brain slices were treated with 0.5 mg/mL lysolecithin (Sigma-Aldrich) at 3 days post isolation for 16 h. Next, brain slices were allowed to recover in culture medium for 1 day and subsequently treated daily with vehicle or BMDM-derived EVs (2 × 10^9^ EVs/mL) for 1 week, followed by histological and biochemical analysis.

### OPC isolation and differentiation

OPCs were isolated as described previously (Dierckx et al., 2022). Briefly, OPCs are obtained from P0-2 wild-type C57BL/6J mice cerebral cortices, and cells were enzymatically dissociated for 20 min at 37°C with papain (3 U/mL, Sigma-Aldrich), diluted in Dulbecco’s Modified Eagle Medium (DMEM; Gibco) and supplemented with 1 mM L-cysteine (Sigma-Aldrich), and DNase I (20 µg/mL, Sigma-Aldrich). The resulting mixed glial cells were cultured in DMEM supplemented with 10% FCS, 50 U/mL penicillin, and 50 U/mL streptomycin in poly-L-lysine (PLL; 50 µg/mL, Sigma-Aldrich)-coated T75 culture flasks at 37°C and 8.5% CO_2_. Medium change was performed on day 4, 7, and 11, and from day 7, cultures were supplemented with bovine insulin (5 µg/mL, Sigma-Aldrich). On day 14, OPCs were isolated using the orbital shake-off method. Briefly, cultures were first shaken at 75 rpm and 37°C for 45 min to detach microglia, and next, cultures were shaken for 16 h at 250 rpm and 37°C to detach OPCs. Subsequently, OPC-containing supernatant was incubated on a Petri dish for 20 min at 37°C and centrifuged at 300 g. Afterwards, OPCs were plated on PLL-coated wells at a density of 3 × 10^5^ cells/mL in proliferation medium (DMEM medium supplemented with 0.3 mM transferrin, 0.1 mM putrescin, 0.02 mM progesterone, 0.2 µM sodium selenite, 0.5 µM triiodothyronine, 0.5 mM L-thyroxin, 0.8 mM bovine insulin, 50 U/mL penicillin, 50 U/mL streptomycin, 2% horse serum, 2% B27 supplement, 5 ng/µL platelet-derived growth factor α (PDGFα), and 5 ng/µL fibroblast growth factor (FGF); all from Sigma-Aldrich except for penicillin/streptomycin (Invitrogen), B27 (in house production as described by Chen et al.), and PDGFα/FGF (Peprotech) for 2 days. Next, OPCs were cultured in differentiation medium (proliferation medium without PDGFα/FGF) and treated daily for 6 days with vehicle, conditioned medium, or BMDM-derived EVs (4 × 10^8^ EVs/mL).

### Transmission electron microscopy

EVs were negatively stained by 2% uranyl acetate. Briefly, 10 µL of EVs at an average concentration of 10^11^ EVs/mL were drop-casted on copper grids (TEDPELLA, #01824) for 60 s and dried with filter paper. Next, samples were stained by two steps of 3 µL 2% uranyl acetate for 30 s. Mouse brain samples were fixed with 2% glutaraldehyde. Next, post-fixation was done with 2% osmiumtetroxide in 0.05 M sodium cacodylate buffer for 1 h at 4°C. Dehydration of the samples was performed by ascending concentrations of acetone. Next, the dehydrated samples were impregnated overnight in a 1:1 mixture of acetone and araldite epoxy resin. Next, the samples were embedded in araldite epoxy resin at 60°C and were cut in slices of 70 nm, perpendicular to the corpus callosum, with a Leica EM UC6 microtome. The slices were transferred to copper grids (Veco B.V) that were coated with 0.7% formvar. Analysis was performed with a Jeol JEM-1400 Flash at 80 kV equipped with an Emsis 20 NP XAROSA camera system; for the EV images were collected images in at least 10 different grid regions and for the mouse brain sections around 5-12 images were taken. ImageJ was used to calculate the g-ratio (ratio of the inner axonal diameter to the total outer diameter). All analyses were conducted by observers blinded to the experimental arm of the study.

### Immunofluorescence

Murine OPCs were cultured on PLL-coated glass cover slides and fixed in 4% paraformaldehyde (PFA) for 20 min at room temperature. Cerebellar brain slices were fixed in 4% PFA for 15 min at room temperature. Brain tissue of cuprizone mice was isolated, snap-frozen, and sectioned with a Leica CM1900UV cryostat (Leica Microsystems) to obtain 10 µm slices. Cryosections were fixed in ice-cold acetone for 10 min at -20°C. Immunostaining and analysis of cryosections were performed as described previously (Bogie et al., 2017). ImageJ was used to align the corpus callosum, followed by determination of the MBP, Olig2, and CC1 positive signal in this area. To stain OPCs, samples were incubated with the relevant primary antibodies diluted in blocking buffer (1x PBS + 1% BSA + 0.1% Tween-20). To stain cerebellar brain slices, samples were incubated with relevant primary antibodies diluted in blocking buffer (1x PBS + 5% horse serum + 0.3% Triton X-100). Analysis of OPCs and cerebellar brain slices was performed using the LeicaDM2000 LED fluorescence microscope and the LSM 880 confocal microscope (Zeiss), respectively. ImageJ was used to determine the MBP/O4 positive area per cell and to define the Sholl analysis parameters related to process complexity and branching (sum intersections, average intersections/Sholl ring, end radius) in OPCs as previously described (Murtie et al., 2007), and to define the Olig2 and CC1 positive cells in the cerebellar brain slices, which is presented as the percentage Olig2^+^ CC1^+^ cells/CC1^+^ cells, and the myelination index (MBP^+^ NF^+^ axons/NF^+^ axons) which is presented in a relative normalized manner. Three-dimensional analysis of cerebellar brain slices was performed using the z-stack feature, and images were 3D rendered using the 3D rendering software vaa3d (Peng et al., 2014). Representative images shown in figures are digitally enhanced to increase the readability. All analyses were conducted by observers blinded to the experimental arm of the study.

### LC-ESI-MS/MS

EV pellets were diluted in 700 µL 1x PBS with 800 µl 1 N HCl:CH_3_OH 1:8 (v/v), 900 µL ChCl_3_ and 200 µg/mL of the antioxidant 2,6-di-tert-butyl-4-methylphenol (BHT; Sigma-Aldrich). 3 µL of SPLASH LIPIDOMIX Mass Spec Standard (Avanti Polar Lipids) was added to the extract mix. The organic fraction was evaporated at room temperature using the Savant Speedvac spd111v (Thermo Fisher), and the remaining lipid pellet was reconstituted in 100% ethanol. Lipid species were analyzed by liquid chromatography-electrospray ionization tandem mass spectrometry (LC-ESI-MS/MS) on a Nexera X2 UHPLC system (Shimadzu) coupled with hybrid triple quadrupole/linear ion trap mass spectrometer (6500+ QTRAP system; AB SCIEX). Chromatographic separation was performed on a XBridge amide column (150 mm x 4.6 mm, 3×5 µm; Waters) maintained at 35°C using mobile phase A [1 mM ammonium acetate in water-acetonitrile 5:95 (v/v)] and mobile phase B [1 mM ammonium acetate in water-acetonitrile 50:50 (v/v)] in the following gradient: (0-6 min: 0% B → 6% B; 6-10 min: 6% B → 25% B; 10-11 min: 25% B → 98% B; 11-13 min: 98% B → 100% B; 13-19 min: 100% B; 19-24 min: 0% B) at a flow rate of 0.7 mL/min which was increased to 1.5 mL/min from 13 minutes onwards. Sphingomyelin (SM) and cholesteryl esters (CE) were measured in positive ion mode with a precursor scan of 184.1 and 369.4. Triglycerides (TG), diglycerides, and monoglycerides were measured in positive ion mode with a neutral loss scan for one of the fatty acyl moieties. Phosphatidylcholine (PC), phosphatidylethanolamine (PE), phosphatidylglycerol (PG), phosphatidylinositides (PI), and phosphatidylserines (PS) were measured in negative ion mode by fatty acyl fragment ions. Lipid quantification was performed by scheduled multiple reactions monitoring, the transitions being based on the neutral losses or the typical product ions as described above. The instrument parameters were as follows: Curtain Gas = 35 psi; Collision Gas = 8 a.u. (medium); IonSpray Voltage = 5500 V and -4500 V; Temperature = 550°C; Ion Source Gas 1 = 50 psi; Ion Source Gas 2 = 60 psi; Declustering Potential = 60 V and -80 V; Entrance Potential = 10 V and -10 V; Collision Cell Exit Potential = 15 V and -15 V. Peak integration was performed with the Multiquant TM software version 3.0.3. Lipid species signals were corrected for isotopic contributions (calculated with Python Molmass 2019.1.1) and were quantified based on internal standard signals and adhere to the guidelines of the Lipidomics Standards Initiative. Only the detectable lipid classes and fatty acyl moieties are reported in this manuscript.

### Cholesterol measurements

Cholesterol levels of BMDM-derived EVs were defined by using the Amplex Red Cholesterol Assay kit (Thermo Fisher) according to the manufacturer’s instructions. Fluorescence was measured using the CLARIOstar PLUS microplate reader (BMG Labtech).

### Cholesterol depletion/repletion

IL-4-stimulated BMDM-derived EVs were depleted of cholesterol by incubation with 2.5% and 5% m/v MβCD for 1 h at 37°C. LPS-stimulated BMDM-derived EVs were repleted with cholesterol by incubating the EVs with 2.5% and 5% m/v MβCD/cholesterol complexes (molar mass 8:1) overnight at 37°C while shaking at 250 rpm. After cholesterol depletion/repletion, unincorporated MβCD and MβCD:cholesterol complexes were removed by means of SEC, and EVs were concentrated using Amicon 0.5mL 10 kDa filters. Depletion/repletion of cholesterol was verified using the Amplex Red Cholesterol Assay kit.

### EV uptake

EVs were fluorescently labelled with 1,1’-dioctadecyl-3,3,3’,3’-tetramethylindo-carbocyanine perchlorate (DiI; Sigma-Aldrich) for 30 min at 37°C. Next, unincorporated DiI was removed using SEC and samples were concentrated using the Amicon Ultra 0.5 mL 10 kDa filter. To investigate the effect of inhibiting specific EV uptake pathways, OPCs, either cultured on coverslips or not, were pre-incubated with Cytochalasin D, chlorpromazine hydrochloride, nystatin, imipramine hydrochloride, wortmannin, omeprazole and MβCD for 30 mins. Next, OPCs were exposed to 4 × 10^8^ DiI-labelled EVs/mL for 3 h and subsequently washed with PBS to remove unincorporated DiI-labelled EVs. Thereafter, OPCs cultured on coverslips were fixated with 4% PFA for 20 min at room temperature, counterstained with DAPI (Sigma-Aldrich), imaged using the Leica DM2000 LED fluorescence microscope (Leica Microsystems), and analyzed using ImageJ. Similarly, OPCs not cultured on coverslips were collected and analyzed for fluorescence intensity using the FACS Calibur (BD Biosciences).

### Luciferase-based nuclear receptor reporter assay

To determine the activation of LXRα and LXRβ, luciferase-based reporter assays were performed using the ONE-GloTM Luciferase Assay System kit (Promega). COS7 cells were transfected with bacterial plasmid constructs expressing luciferase under the control of the ligand-binding domain for LXRα or LXRβ, which were kindly provided by Prof. dr. Bart Staels (Université de Lille, INSERM, France). Cells were grown to 60% confluency in 60 mm petri dishes and subsequently transfected with 1.8 µg plasmid DNA, including 0.2 µg pGAL4hLXRα or pGAL4hLXRβ, 1 µg pG5-TK-GL3, and 0.6 µg pCMV-β-galactosidase. JetPEI (Polysciences) was used as transfection reagent. After overnight incubation, transfected cells were seeded at a density of 10,000 cells/well in a 96-well plate in DMEM medium enriched with 50 u/mL penicillin and 50 U/mL streptomycin, and treated with vehicle, BMDM-derived EVs, or T0901317 for 24 h. Following treatment, cells were lysed in lysis buffer (25 mM Glycyl-Glycine, 15 mM MgSO_4_, 4 mM EGTA, and 1x Triton, all from Sigma-Aldrich). To correct for transfection efficacy, β-galactosidase activity was measured using cell lysate (10%) in β-gal buffer, consisting of 20% 2-Nitrophenyl β-D-galactopyranoside (ONGP; Sigma-Aldrich) and 80% Buffer-Z (0.1 M Na_2_HPO_4_, 10 mM KCl, 1 mM MgSO_4_, and 3.4 µL/mL 2-mercaptoethanol; all from Sigma-Aldrich). Luminescence and absorbance (410 nm) were measured using the CLARIOstar PLUS microplate reader.

### Statistical analysis

Data were statistically analyzed using GraphPad Prism v8 and are reported as mean ± SEM. The D’Agostino and Pearson omnibus normality test was used to test for normal distribution. When datasets were normally distributed, an ANOVA (Tukey’s post hoc analysis) or two-tailed unpaired Student’s *t*-test (with Welch’s correction if necessary) was used to determine statistical significance between groups. If datasets did not pass normality, the Kruskal-Wallis or Mann-Whitney analysis was applied. P values < 0.05 were considered to indicate a significant difference (*, p < 0.05; **, p < 0.01; ***, p < 0.001).

## Results

### Extracellular vesicles released by RAMs improve OPC differentiation

Macrophages display tremendous phenotypical plasticity in neurological disorders, a reflection of dynamic intracellular and environmental changes (Bogie et al., 2020). To date, however, the molecular mechanisms and factors that drive the harmful and benign effector functions of macrophages in CNS disorders remain poorly understood. By exposing OPC cultures to conditioned medium of naive macrophages (M0), DAMs, or RAMs, we show that the secretome underlies the regenerative impact of RAMs on OPC maturation. Specifically, fluorescent staining demonstrated an increased ratio of the mature OLN marker MBP over the pre-OLN marker O4 in OPC cultures exposed to conditioned medium of RAMs (Fig. 1A-C). To confirm enhanced differentiation of OPCs at a morphological level, Sholl analysis was performed. Here, OPCs exposed to the secretome of RAMs displayed an increased number of dendrite branches as well as a more complex dendrite branch geometry (Fig. 1D-E), morphological characteristics typically associated with enhanced OLN maturation (Livesey et al., 2016). Surprisingly, conditioned medium of DAMs did not impact the MBP/O4 ratio and morphological parameters in the Sholl analysis (Fig. 1B-E), nor did the dendrite ending radius differ in OPCs exposed to the secretome of naive macrophages, DAMs, and RAMs (Fig. 1F). Altogether, these findings show that RAM-secreted soluble factors, at least partially, enhance OPC differentiation *in vitro*.

**Figure 1.**
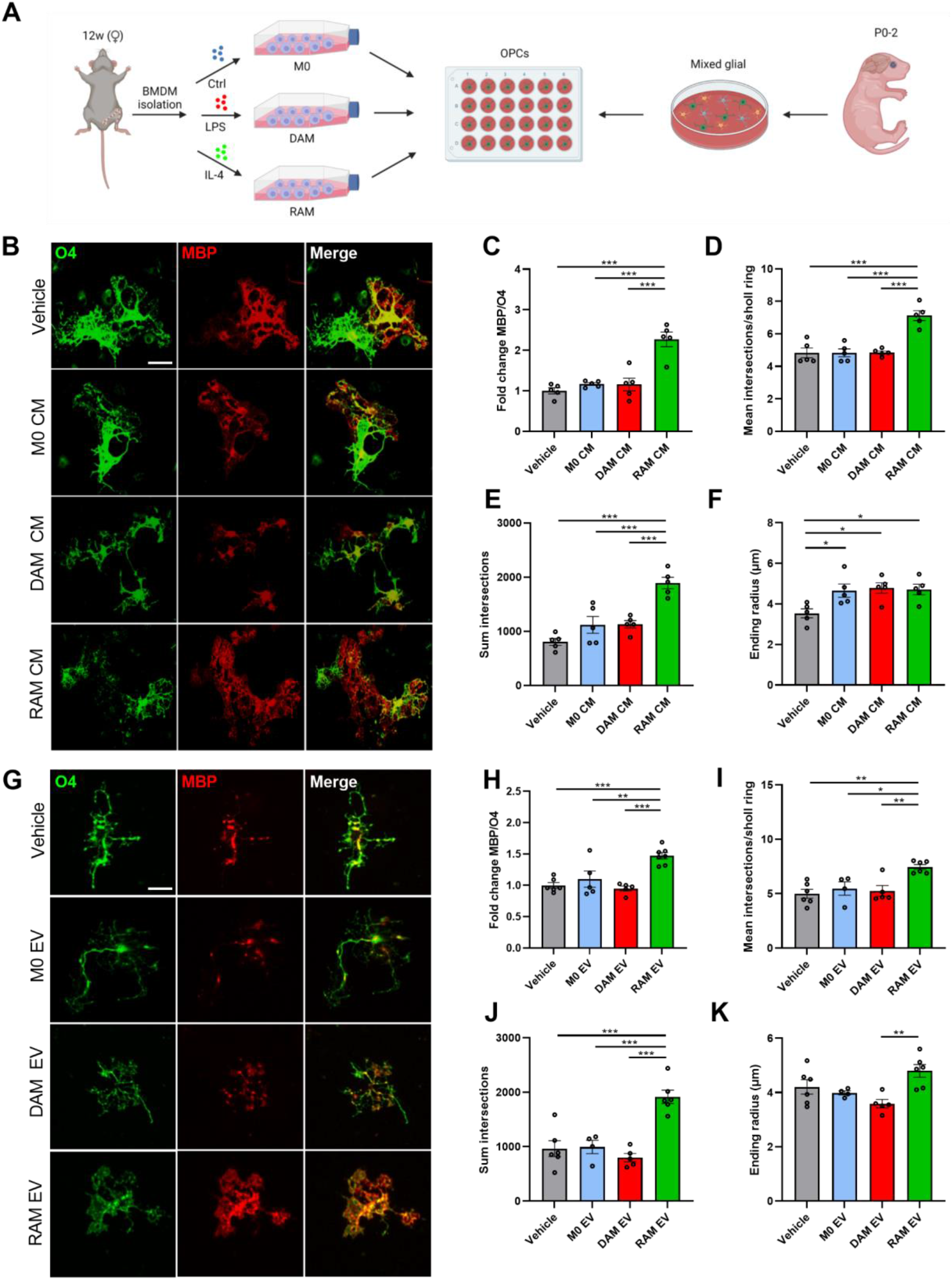
Extracellular vesicles released by RAMs improve OPC differentiation. **A**) Schematic representation showing the isolation of oligodendrocyte precursor cells (OPCs) and their stimulation with conditioned medium (CM) and extracellular vesicles (EVs) released by naive macrophages (M0, PBS-treated), disease-associated macrophages (DAMs, LPS-stimulated), and repair-associated macrophages (RAMs, IL-4-stimulated). Created with biorender.com. **B-K**) Representative immunofluorescent images (B,G) and quantification (C-F, H-K) of OPC maturation exposed to vehicle (BMDM culture medium (B-F), PBS (G-K)), conditioned medium (B-F) or EVs (G-K) isolated from M0, DAMs, and RAMs for 6 days. OPCs were stained for MBP (mature oligodendrocyte) and O4 (premature oligodendrocyte), and differentiation was quantified by measuring the MBP/O4 ratio (C, H) or applying the Sholl analysis (mean interactions/ring, sum intersections, and ending radius; D-F, I-K). OPCs were treated with conditioned medium produced by 1.5 × 10^5^ BMDMs or 4 × 10^8^ EVs/mL. Scale bar, 25 µm. Results are pooled from or representative of four biological replicates (n = 4-6 cultures). Data are represented as mean ± SEM and statistically analyzed using the one-way ANOVA test followed by Tukey’s multiple comparison test. *, p < 0.05; **, p < 0.01; ***, p < 0.001.

Emerging evidence points towards the pathophysiological significance of macrophage-derived EVs in the initiation and resolution of chronic inflammatory disorders (Jiang et al., 2022; Liu et al., 2021; Long et al., 2021). To define whether EVs underlie the reparative impact of RAMs on OPC maturation (Fig. 1), OPCs were exposed to EVs isolated from naive macrophages, DAMs, or RAMs (experimental design in Suppl. Fig. 1A). First, EVs were characterized according to the “Minimal Information for Studies of Extracellular Vesicles (MISEV)” guidelines (Witwer et al., 2021). Transmission electron microscopy demonstrated EVs displaying the typical spherical and cup-shape morphology (Suppl. Fig. 1B). Nanoparticle Tracking Analysis (NTA) further established similar distribution curves of EVs released by naive macrophages, DAMs, and RAMs, with the majority of EVs displaying a particle size ranging from 50 to 150 nm (Suppl. Fig. 1B). Finally, immunoblotting showed that all EV isolates were positive for CD81 and Flottilin1, while lacking the endoplasmic marker GRP94 (Suppl. Fig. 1C). Collectively, these findings confirm the isolation of small EVs. Having established the efficient isolation of EVs from macrophages, we next assessed their impact on OPC differentiation. Mirroring the pro-regenerative impact of the RAM secretome, fluorescent staining demonstrated that EVs released by RAMs enhanced OPC maturation, evidenced by an increased MBP/O4 ratio (Fig. 1G-H). Consistent with enhanced OPC maturation, Sholl analysis demonstrated higher dendrite branch numbers and complexity in OPCs exposed to EVs released by RAMs and revealed a higher dendrite ending radius in EVs released by RAMs compared to DAM-derived EVs (Fig. 1I-K). EVs released by naive macrophages and DAMs did not impact OPC maturation (Fig. 1G-H), nor did dendrite ending radius differ in OPCs exposed to EVs released by naive macrophages and RAMs (Fig. 1I-K). In aggregate, these findings indicate that RAMs promote the maturation of OPCs *in vitro*, at least partially, through the release of EVs.

### Extracellular vesicles released by RAMs improve remyelination

Given that EVs released by RAMs enhanced OPC differentiation, we next sought to define their impact on remyelination. Remyelination was first studied using *ex vivo* cerebellar brain slices. To this end, demyelinated brain slices were exposed to EVs isolated from naive macrophages, DAMs, and RAMs (experimental design in Fig. 2A). Fluorescent staining demonstrated increased colocalization of myelin (MBP) and axons (neurofilament, NF) in brain slices treated with RAM-derived EVs, indicating enhanced axonal remyelination (Fig. 2B-C). Improved remyelination was confirmed using three-dimensional reconstructions (Fig. 2B). Akin to our *in vitro* findings, RAM-derived EVs increased the percentage of CC1^+^ mature OLNs within Olig2^+^ oligodendroglial lineage cells (Fig. 2D-E). Remyelination efficiency and the percentage of CC1^+^ Olig2^+^ cells were not affected in brain slices exposed to EVs released by DAMs as well as naive macrophages (Fig. 2B-E), similar to our *in vitro* OPC cultures. These findings indicate that EVs released by RAMs promote remyelination in *ex vivo* cerebellar brain slices, and support the notion that enhanced maturation of OPCs underlies their reparative impact.

**Figure 2:**
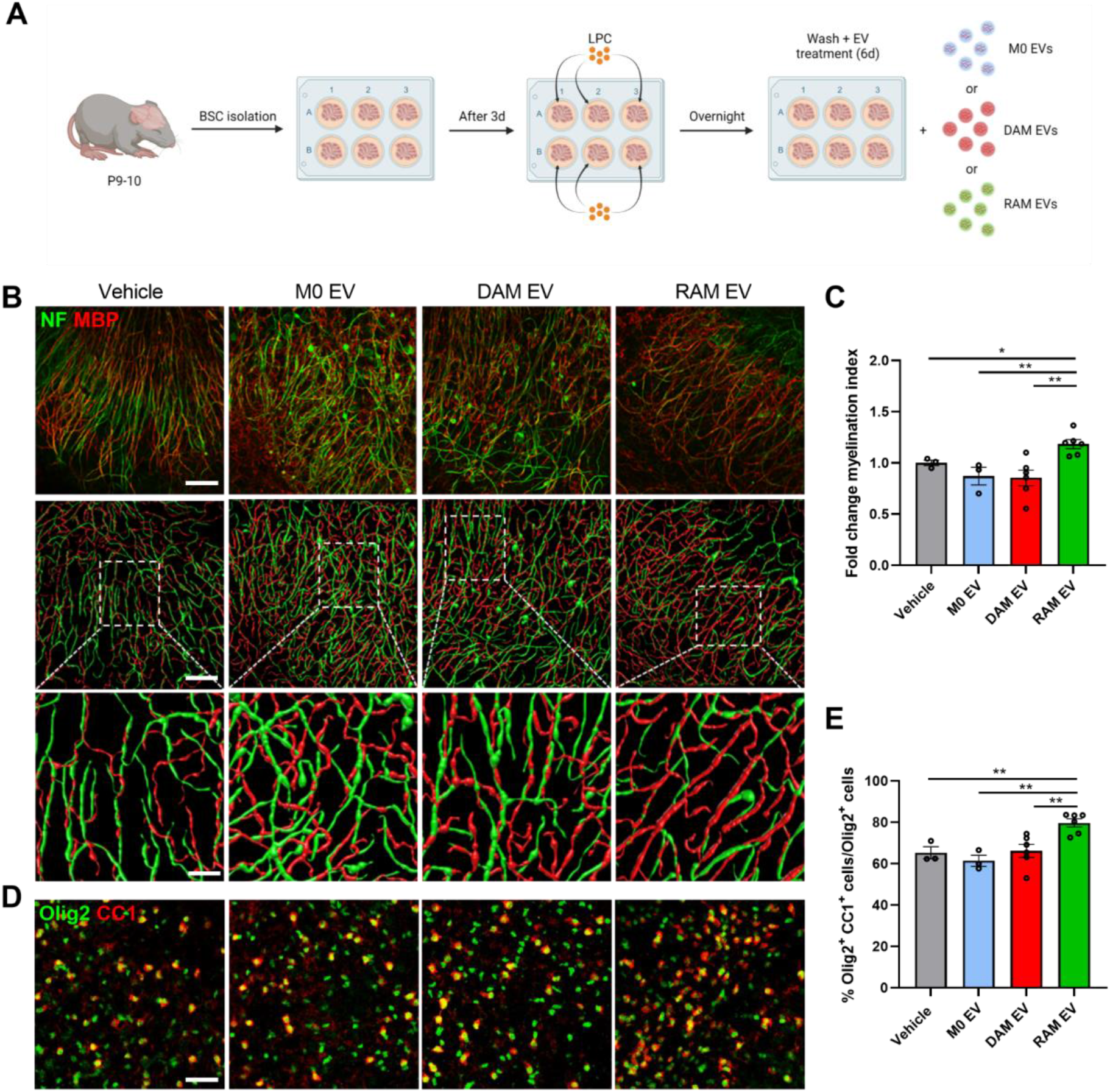
Extracellular vesicles released by RAMs improve remyelination in cerebellar brain slices. **A**) Schematic representation showing the isolation and culture of cerebellar brain slices as well as their stimulation with vehicle (PBS), or extracellular vesicles (EVs) released by naive macrophages (M0, PBS-treated), disease-associated macrophages (DAMs, LPS-stimulated), and repair-associated macrophages (RAMs, IL-4-stimulated). LPC = lysolecithin, demyelinating compound. Created with biorender.com. **B**,**D**) Representative images and three-dimensional reconstruction of immunofluorescent MBP/NF (B) and Olig2/CC1 (D) stains of cerebellar brain slices treated with vehicle or EVs isolated from M0, DAMs, and RAMs. Slices were treated with 4 × 10^9^ EVs/mL. Scale bars, 100 µm (B, (row 1, 2) D); 25 µm (B, row 3). **C**) Relative number of MBP^+^ NF^+^ axons out of total NF^+^ axons in cerebellar brain slices treated with vehicle or EVs released by M0, DAMs, and RAMs (n = 3-6 slices). **E**) Percentage Olig2^+^ CC1^+^ cells within the Olig2^+^ cell population in cerebellar brain slices treated with vehicle or EVs released by M0, DAMs, and RAMs (n = 3-6 slices). Results are pooled from or representative of three independent experiments. Data are represented as mean ± SEM and statistically analyzed using the Kruskal-Wallis test followed by Dunn’s multiple comparison test. *, p < 0.05; **, p < 0.01.

To evaluate the significance of these findings *in vivo*, the cuprizone-induced de- and remyelination model was used. Cuprizone feeding leads to reproducible toxic demyelination in distinct brain regions, the corpus callosum (CC) in particular. Upon cessation of cuprizone feeding, spontaneous remyelination occurs, as evidenced by an increased MBP levels, decreased g-ratio (the ratio of the inner axonal diameter to the total outer diameter), and higher percentage of CC1^+^ Olig2^+^ cells in the CC of mice 1 week after cessation of cuprizone feeding (5w+1) (Suppl. Fig. 2A-D). A single intracerebroventricular administration after demyelination demonstrated that EVs released by RAMs enhanced MBP abundance in the CC of cuprizone animals as compared to animals exposed to vehicle or EVs released by naive macrophages and DAMs (Fig. 3A-C). Accordingly, transmission electron microscopy showed a decreased g-ratio and increased percentage of myelinated axons in the corpus callosum of mice treated with EVs released by RAMs (Fig. 3B, D-E). In particular, small diameter axons showed thicker myelin sheaths in mice that were treated with RAM-derived EVs (Suppl. Fig. 2E). Moreover, intracerebroventricular administration of RAM-derived EVs increased the number of mature OLNs, evidenced by an increased percentage of CC1^+^ Olig2^+^ cells. Again, EVs released by naive macrophages and DAMs did not affect axonal remyelination and the numbers of CC1^+^ Olig2^+^ cells (Fig. 3B-E). Collectively, these findings indicate that EVs released by RAMs boost remyelination, likely through enhancing the intrinsic capacity of OPCs to mature into myelinating OLNs.

**Figure 3:**
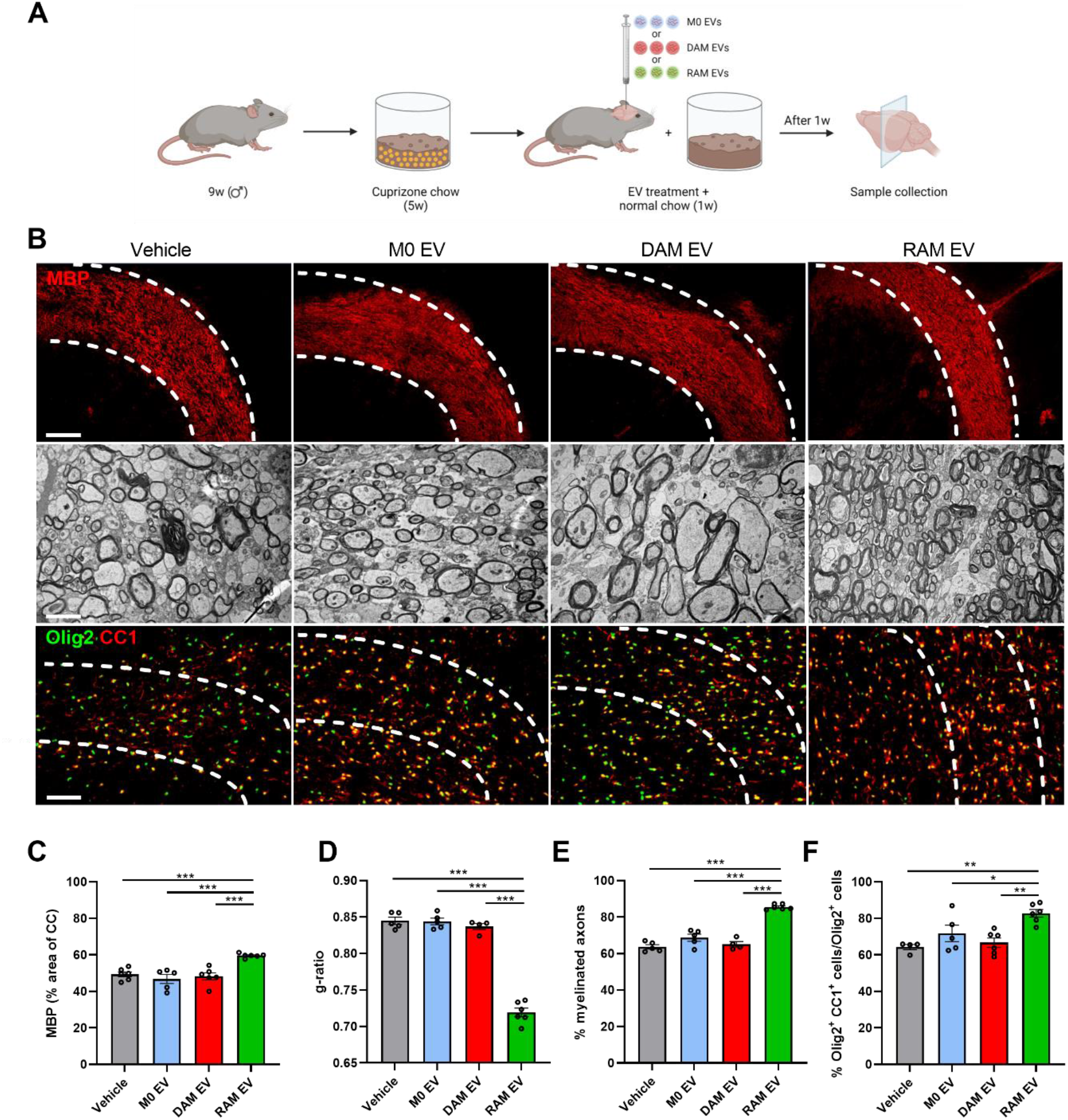
Extracellular vesicles released by RAMs enhance remyelination in the cuprizone model. **A**) Schematic representation showing the experimental pipeline used to assess the impact of extracellular vesicles (EVs) released by naive macrophages (M0, PBS-treated), disease-associated macrophages (DAMs, LPS-stimulated), and repair-associated macrophages (RAMs, IL-4-stimulated) on remyelination in the cuprizone model. Created with biorender.com. **B**) Representative images of immunofluorescent MBP and Olig2/CC1 stains and transmission electron microscopy analysis of the corpus callosum (CC) from mice treated intracerebroventricularly with vehicle (PBS) or EVs released by M0, DAMs, and RAMs. The outer border of the CC is demarcated by the dotted line. Mice were injected with vehicle or EVs (1 × 10^9^ EVs) after demyelination (5w), and analysis was performed during remyelination (5w+1). Scale bar, 200 µm (rows 1, 3) and 2 µm (row 2). **C**) Quantification of the MBP^+^ area of the CC from cuprizone mice treated with vehicle or EVs during remyelination (5w+1) (n = 5-6 animals, 3 images/animal). **D-E**) Analysis of the g-ratio (the ratio of the inner axonal diameter to the total outer diameter, D), and percentage myelinated axons (E) in CC from cuprizone mice treated with vehicle or EVs during remyelination (5w+1) (n = 3-5 animals, 3 images/animal, 100-150 axons/image). **F**) Quantification of the percentage Olig2^+^ CC1^+^ cells out of total Olig2^+^ cells in the CC of cuprizone animals treated with vehicle or EVs during remyelination (5w+1) (n = 4-6 animals, 3 images/animal). Data are represented as mean ± SEM and statistically analyzed using the Kruskal-Wallis test followed by Dunn’s multiple comparison test. *, p < 0.05; **, p < 0.01; ***, p < 0.001.

### Cholesterol abundance controls the impact of EVs released by macrophages on OPC maturation

Ample evidence indicates that OPC differentiation and remyelination cause a surge in lipid demand (Dimas et al., 2019; Hubler et al., 2018; Saher et al., 2005). Given that 1) EVs are carriers of lipids, 2) the importance of immunometabolism in driving macrophage effectors functions, and 3) EV-associated lipids being acknowledged to affect neuropathological processes (Bogie et al., 2020; Goossens et al., 2019; Grey et al., 2015; Krämer-Albers et al., 2007; Oishi et al., 2017; Vanherle et al., 2020; Wang et al., 2012), we next assessed the impact of EV-associated lipids on remyelination. Here, we provide evidence that the lipid fraction of EVs released by RAMs markedly enhanced the MBP/O4 ratio as well as dendrite branch numbers and complexity of OPCs (Fig. 4A-D; Suppl. Fig. 3A). Similar to our previous findings, lipids within EVs released by naive macrophages and DAMs did not impact OPC maturation. To identify lipids involved in driving the regenerative impact of RAM-derived EVs, electrospray ionization tandem mass spectrometry (ESI-MS/MS) analysis was performed on EVs. Interestingly, while the vast majority of glycerolipids, glycerophospholipids, and sphingolipids were unaltered in all EV subsets, EVs released by RAMs showed a marked increase in cholesteryl ester (CE) levels (Fig. 4E). In-depth profiling did not identify changes in the relative abundance (% of class) of divergent fatty acid species within CE of RAM- and DAM-derived EVs, indicating an overall increase of all CE species (Fig. 4F). Increased levels of CE, as well as free and total cholesterol, were confirmed using the Amplex Red Cholesterol Assay (Fig. 4G). To establish whether increased cholesterol abundance underpins the regenerative impact of EVs released by RAMs, a cholesterol-depletion and -enrichment strategy using unloaded and cholesterol-loaded MβCD was applied (Experimental design in Fig. 4H and I). MβCD efficiently depleted and enriched RAMs and DAMs of cholesterol, respectively (Suppl. Fig. 3B, F). As expected, EVs depleted of cholesterol showed a reduced particle size, while the size of those enriched in cholesterol was increased (Suppl. Fig. 3C, G). Given that high levels of MβCD (5%) noticeably decreased the number of EVs, potentially due to cholesterol-associated changes in membrane integrity, we opted to continue with EVs modified with 2.5% MβCD and cholesterol-loaded MβCD (Suppl. Fig. 3D, H). Notably, cholesterol depletion and enrichment did not affect the uptake of EVs by OPCs (Suppl. Fig. 3E, I). In support of the importance of EV-associated cholesterol in driving OPC maturation, the pro-regenerative impact of EVs released by RAMs on OPC maturation and morphology was nullified upon cholesterol depletion (Fig. 4J-M, Suppl. Fig. 3H). Vice versa, EVs released by DAMs gained the ability to promote OPC maturation once enriched with cholesterol, with values reaching those observed in OPC cultures exposed to non-modified EVs released by RAMs (Fig. 4J-M, Suppl. Fig. 3H). Altogether, these findings highlight that EVs are essential for cholesterol trafficking and that changes in cholesterol abundance dictate, at least partially, the pro-regenerative impact of EVs released by macrophages on OPC maturation.

**Figure 4:**
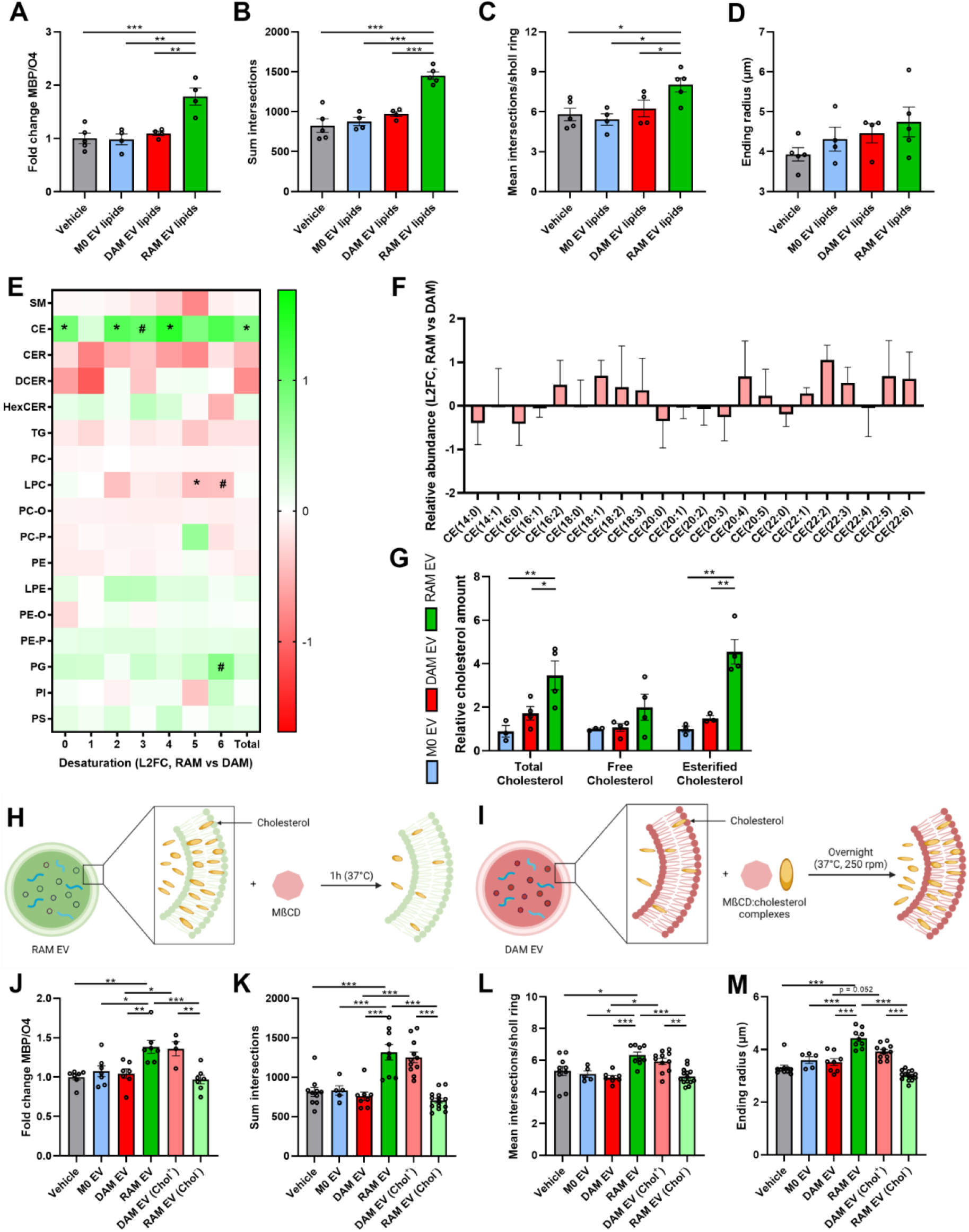
Cholesterol abundance controls the impact of EVs released by macrophages on OPC maturation. **A-D**) Quantification of OPC maturation following 6 d exposure to lipids isolated from EVs released by naive macrophages (M0, PBS-treated), disease-associated macrophages (DAMs, LPS-stimulated), and repair-associated macrophages (RAMs, IL-4-stimulated). OPCs were stained for MBP (mature oligodendrocyte) and O4 (premature oligodendrocyte), and differentiation was quantified by measuring the MBP/O4 ratio (A) or applying the Sholl analysis (mean interactions/ring, sum intersection, and ending radius; B-D). OPCs were treated with 4 × 10^8^ EVs/mL (n = 4-5 cultures). **E**) Liquid chromatography electrospray tandem mass spectrometry (LC-ESI-MS/MS) analysis to measure lipid species of EVs released by DAMs and RAMs (n = 3 isolates). Data are depicted as log^2^ fold change (L2FC). SM; sphingomyelin, CE; cholesterol esters, TG; triglycerides, (D/H)CER; (dihydro/hexosyl)ceramide, (L)PC(-O/-P); (lyso,-alkyl/-alkenyl) phosphatidylcholine, (L)PE(-O/-P); (lyso, -alkyl/-alkenyl) phosphatidylethanolamine, PG; phosphatidylglycerol, and PI; phosphatidylinositol. **F**) Relative abundance of fatty acyl moieties within CE group is shown (RAM EVs vs. DAM EVs). Data are depicted as L2FC (n = 3 isolates). **G**) Quantification of total cholesterol, free cholesterol, and esterified cholesterol in EVs isolated from M0, DAMs, and RAMs (n = 3-4 isolates). **H-I**) Schematic representation of the experimental pipeline used for cholesterol depletion (H) and enrichment (I) of EVs released by RAMs and DAMs, respectively. Created with biorender.com. **J-M**) Quantification of OPC maturation following 6 d exposure to EVs isolated from M0, DAMs, and RAMs, as well as EVs isolated from DAMs and RAMs that were enriched (Chol^+^) with or depleted (Chol^-^) of cholesterol, respectively. OPCs were stained for MBP (mature oligodendrocyte) and O4 (premature oligodendrocyte), and differentiation was quantified by measuring the MBP/O4 ratio (J) or applying the Sholl analysis (mean interactions/ring, sum intersection, and ending radius; K-M). OPCs were treated with 4 × 10^8^ EVs/mL (n = 5-12 cultures). All results are pooled from three biological replicates. Data are represented as mean ± SEM and statistically analyzed using the one-way ANOVA test followed by Tukey’s multiple comparison test (A-D, J-M), the Mann-Whitney test (F), and the Kruskal-Wallis test followed by Dunn’s multiple comparison test (G). #, p < 0.1, *, p < 0.05; **, p < 0.01; ***, p < 0.001.

### Cholesterol abundance dictates the impact of EVs released by macrophages on remyelination

To provide evidence that the cholesterol content in macrophage-derived EVs impacts remyelination in a physiological setting, the cerebellar brain slice and cuprizone models were used. Consistent with our *in vitro* findings, we found that EVs released by RAMs lose their reparative features in brain slices once depleted of cholesterol as compared to their non-modified equivalents, evidenced by reduced colocalization of MBP with NF and lower numbers of CC1^+^ Olig2^+^ mature OLNs (Fig. 5A-D). Mirroring these findings, EVs released by DAMs increased axonal remyelination and OLN levels following cholesterol enrichment as compared to their non-modified counterparts (Fig. 5A-D). Similar to brain slice cultures, intracerebroventricular administration of modified and non-modified EVs confirmed the vital role of cholesterol in driving the reparative features of EVs. Specifically, cholesterol depletion of EVs released by RAMs nullified their beneficial impact on remyelination in the CC as evidenced by increased MBP levels, a higher percentage of myelinated axons, and a decreased g-ratio. Vice versa, cholesterol enrichment of DAM-derived EVs increased their beneficial impact on these pathological parameters (Fig. 5E-H, Suppl. Fig. 4A-B). Accordingly, a lower and higher percentage of CC1^+^ Olig2^+^ mature OLNs was observed in the CC of cuprizone mice treated with cholesterol-depleted RAM EVs and -enriched DAM EVs, respectively, as compared to non-modified equivalents (Fig. 5E-H, Suppl. Fig. 4A-B). In both models, neuropathological changes observed after exposure to MβCD-modified EVs matched those seen after treatment with non-modified EVs released by phenotypically divergent macrophage subsets, further emphasizing the importance of cholesterol in dictating the repair-promoting features of macrophage-derived EVs. Collectively, these findings argue that EVs are cholesterol traffickers during remyelination, and that cholesterol abundance underlies the reparative properties of EVs in demyelinated lesions.

**Figure 5:**
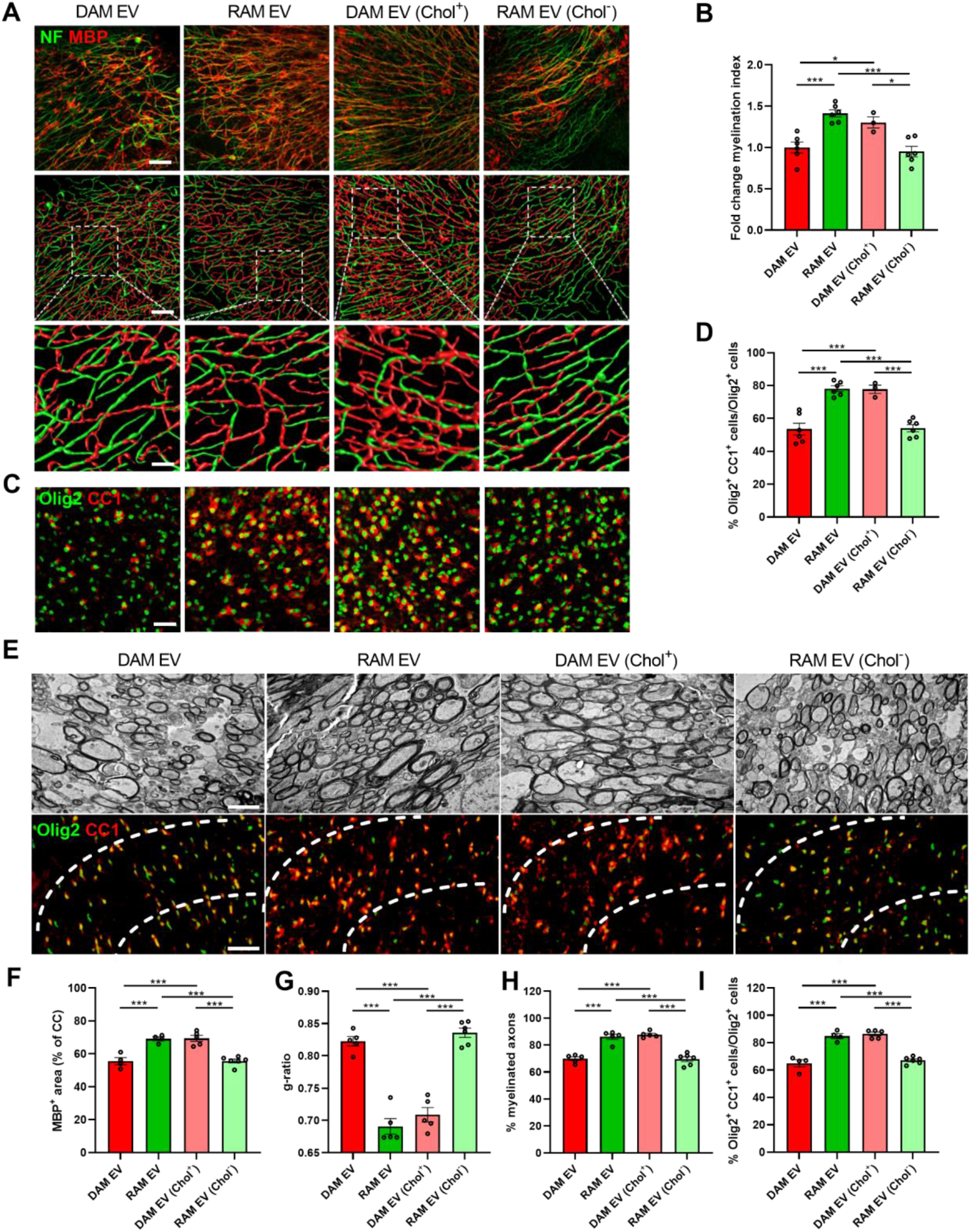
Cholesterol abundance controls the impact of EVs released by macrophages on remyelination. **A-D**) Representative images, three-dimensional reconstruction, and quantification of immunofluorescent MBP/NF (A-B) and Olig2/CC1 stains (C-D) of cerebellar brain slices treated with vehicle or EVs released by disease-associated macrophages (DAMs, LPS-stimulated), and repair-associated macrophages (RAMs, IL-4-stimulated), as well as EVs released by DAMs and RAMs that were enriched (Chol^+^) with or depleted (Chol^-^) of cholesterol, respectively. Relative number of MBP^+^ NF^+^ axons out of total NF^+^ axons (B) and percentage Olig2^+^ CC1^+^ cells within the Olig2^+^ cell population (D) in cerebellar brain slices treated with vehicle or EVs is shown (n = 3-6 slices). Slices were treated with 4 × 10^9^ EVs/mL. Scale bars, 100 µm (A (row 1 and 2), C); 25 µm (A (row 3)). **E**) Representative images of transmission electron microscopy analysis and immunofluorescent Olig2/CC1 stains of the corpus callosum (CC) from mice treated with vehicle or EV subsets. The outer border of the CC is demarcated by the dotted line. Mice were intracerebroventricularly injected with vehicle or EVs (1 × 10^9^ EVs/mL) after demyelination (5w), and analysis was done during remyelination (5w+1). Scale bar, 200 µm (rows 1, 3) and 2 µm (row 2). **F**) Quantification of the MBP^+^ area of the CC from cuprizone mice treated with vehicle or EVs during remyelination (5w+1) (n = 4-6 animals, 3 images/animal). **G-H**) Analysis of the g-ratio (the ratio of the inner axonal diameter to the total outer diameter, G), and percentage of myelinated axons (H) in CC from cuprizone mice treated with vehicle or EVs during remyelination (5w+1) (n = 5-6 animals, 3 images/animal, 100-150 axons/image). **I**) Quantification of the percentage Olig2^+^ CC1^+^ cells out of total Olig2^+^ cells in the CC of cuprizone animals treated with vehicle or EVs during remyelination (5w+1) (n = 4-6 animals, 3 images/animal). Data are represented as mean ± SEM and statistically analyzed using the Kruskal-Wallis test followed by Dunn’s multiple comparison test. *, p < 0.05; **, p < 0.01; ***, p < 0.001.

### Direct membrane fusion of extracellular vesicles provides OPCs with cholesterol required for maturation

EV uptake and incorporation by recipient cells can ensue in several functionally divergent manners that are likely to impact cellular physiology differently (Fig. 6A and (Kalluri and LeBleu, 2020)). To study the molecular mechanisms that underlie the impact of EVs released by RAMs on OPC maturation, we first determined the molecular mechanisms that drive the uptake of EVs by OPCs. By applying pharmacological inhibitors, we found that OPCs incorporate RAM-derived EVs through receptor- and clathrin-mediated endocytosis, lipid rafts, and direct membrane fusion, with inhibition of the latter having the most prominent impact (Fig. 6B). Importantly, pharmacological inhibition of direct fusion counteracted the beneficial impact of RAM EVs on OPC maturation (Fig. 6D-G, Suppl. Fig. 5A). Alongside direct membrane fusion, we and others demonstrated that the formation of cholesterol metabolites is well-established to promote OPC maturation and remyelination by activating the cholesterol-sensing LXRs (Nelissen et al., 2012; Xu et al., 2014). However, by using nuclear receptor luciferase reporter assays, we found that control and modified EVs released by RAMs and DAMs did not enhance LXRα and LXRβ ligation (Suppl. Fig. 5B-C). In aggregate, these results highlight that RAM-derived EVs promote OPC maturation by directly incorporating cholesterol in the membrane of OPCs.

**Figure 6:**
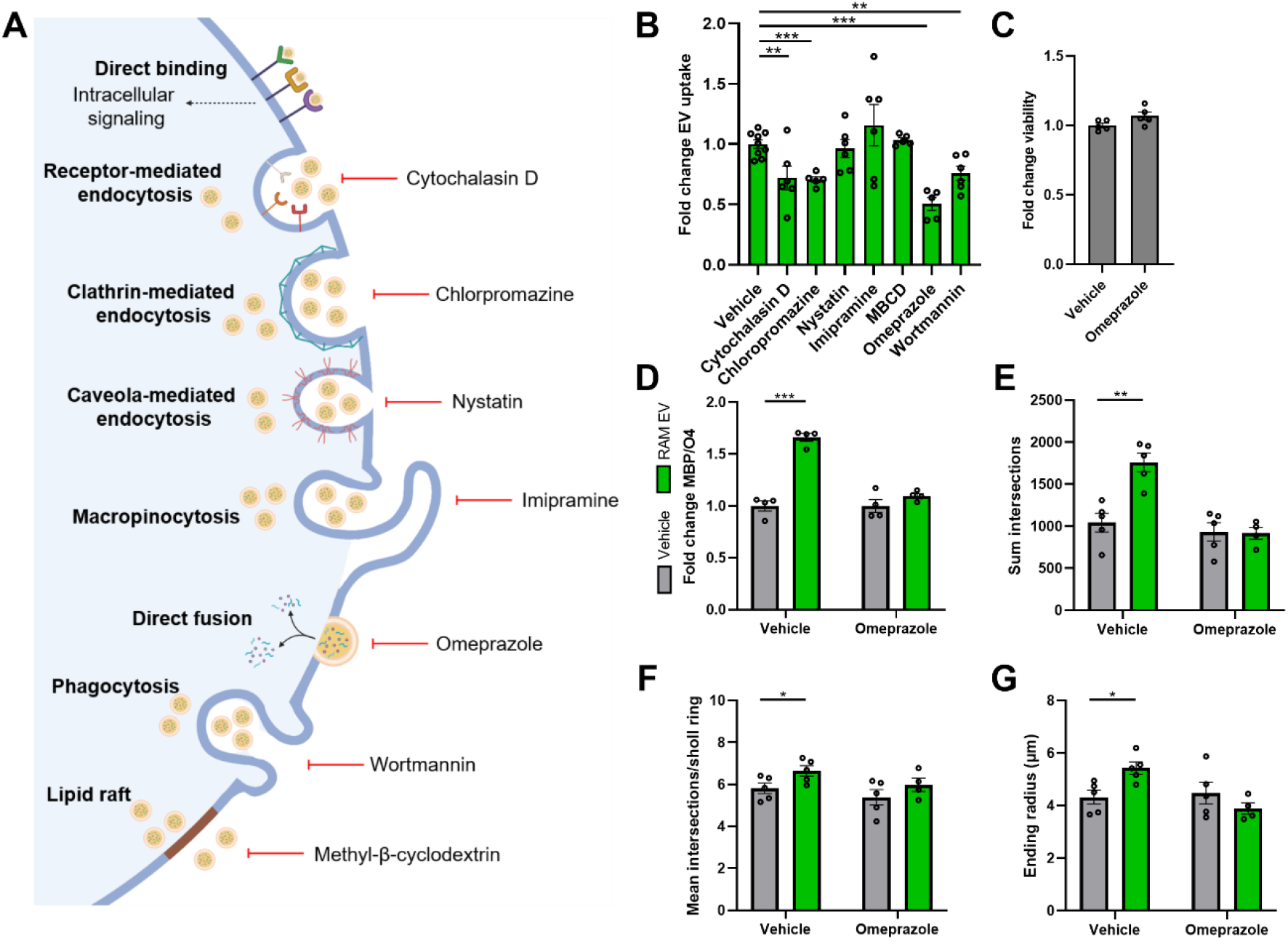
Direct membrane fusion of extracellular vesicles released by RAMs provides OPCs with cholesterol required for maturation. **A**) Schematic representation showing routes of extracellular vesicles (EV) uptake and their associated inhibitors. Created with biorender.com. **B**) Internalization of DiI-labeled EVs released by repair-promoting macrophages (RAMs, IL-4-stimulated) by OPCs pre-treated for 30 min with inhibitors of receptor-mediated endocytosis (cytochalasin D, 10 µM), clathrin-mediated endocytosis (chloropromazine, 50 µM), caveola-mediated endocytosis (nystatin, 25 µM), macropinocytosis (imipramine, 5 µM), direct fusion (omeprazole, 50 µM), phagocytosis (wortmannin, 500 nM), and lipid raft-mediated uptake (methyl-β-cyclodextrin, 1% v/v). OPCs were exposed for 3 h to 4 × 10^8^ EVs/mL (n = 5-9 cultures). **C**) Relative viability of OPCs treated with vehicle or omeprazole for 24 h (n = 5 cultures). **D-G**) Quantification of OPC maturation following 6 d exposure to EVs isolated from RAMs. OPCs were stained for MBP (mature oligodendrocyte) and O4 (premature oligodendrocyte), and differentiation was quantified by measuring the MBP/O4 ratio (D; n = 4 cultures) or applying the Sholl analysis (mean interactions/ring, sum intersection, and ending radius; E-G; n = 5 cultures). OPCs were treated with 4 × 10^8^ EVs/mL. All results are pooled from three biological replicates. Data are represented as mean ± SEM and statistically analyzed using the Kruskal-Wallis test followed by Dunn’s multiple comparison test (B), the Mann-Whitney test (C), and the one-way ANOVA test followed by Tukey’s multiple comparison test (D-G). *, p < 0.05; **, p < 0.01; ***, p < 0.001.

## Discussion

Macrophages display tremendous functional diversity in CNS disorders, being involved in seemingly opposing processes such as neurodegeneration and remyelination (Aydınlı et al., 2022; Bogie et al., 2020; Marschallinger et al., 2020; Miron et al., 2013). Here, we report that RAMs promote OPC maturation and improve remyelination through the release of EVs. Lipidomic analysis revealed that EVs released by RAMs contained higher cholesterol levels, a steroid whose importance in remyelination has been well-established (Mathews et al., 2014; Saher et al., 2005). Cholesterol depletion counteracted the benign impact of EVs released by RAMs on OPC maturation and remyelination. Accordingly, cholesterol enrichment rendered DAM-derived EVs reparative, stipulating the crucial role of EV-associated cholesterol in remyelination, which we found to be directly incorporated into the membrane of OPCs via membrane fusion. Collectively, our findings show that EVs act as cholesterol traffickers in the CNS and that cholesterol abundance underlies their pro-remyelinating properties in demyelinated lesions.

We provide evidence that RAMs promote OPC differentiation *in vitro* and enhance remyelination *ex vivo* and *in vivo* through the release of soluble factors and EVs. Consistent with our findings, other studies defined that phagocytes can release soluble factors that promote CNS remyelination (Boyles et al., 1989; Butovsky et al., 2006; Miron et al., 2013; Yuen et al., 2013). Specifically, a phenotypic switch of disease-to repair-associated microglia was found to coincide with elevated levels of activin-A and increased remyelination (Miron et al., 2013). Furthermore, repair-associated microglia induced by IL-4 boost oligodendrogenesis in an autoimmune-induced demyelination model via the production of IGF-I (Butovsky et al., 2006). Ample evidence further indicates that EVs can have regenerative properties in the CNS and periphery (Guo et al., 2020; Herman et al., 2022; Hervera et al., 2018; Lombardi et al., 2019). Of particular interest, Lombardi et al. demonstrated that microglia co-cultured with immunosuppressive mesenchymal stem cells or stimulated with IL-4 release EVs that promote remyelination, either directly by affecting OPC physiology or indirectly by promoting the formation of reparative astrocytes (Lombardi et al., 2019). Our results now extend these findings by showing that disease-resolving peripheral macrophages, despite differing in their ontogeny and function compared to microglia (Ajami et al., 2011; Chrast et al., 2011; Goldmann et al., 2016; Mildner et al., 2007), promote remyelination through the release of EVs as well. Remarkably, unlike disease-associated microglia (Butovsky et al., 2006), soluble factors and EVs released by DAMs did not negatively impact OPC maturation and remyelination. More research is warranted to assess the molecular mechanisms driving this discrepancy, which might well be related to ontogenetic and functional differences between both phagocyte subsets. Further, given that EVs carry other bioactive molecules, including cytokines, growth factors, and miRNAs, which impact CNS homeostasis and repair (Li et al., 2021; Li et al., 2022; Raffaele et al., 2021), a synergistic effect of cholesterol in combination with these factors cannot be excluded. Collectively, our findings indicate that RAMs promote OPC maturation and remyelination at least partially through the release of EVs.

We show that cholesterol abundance in EVs released by macrophages is an essential determinant of their pro-remyelinating impact in the CNS, with higher cholesterol levels more efficiently promoting remyelination. Accordingly, cholesterol is well-known to be the rate-limiting step in myelin biogenesis, and up to 80% of CNS cholesterol can be found in myelin (Björkhem et al., 2010; Dietschy, 2009; Saher et al., 2005). In the context of remyelination, ample evidence further indicates that changes in the level and synthesis of cholesterol and its metabolites markedly impact remyelination efficacy. In line with the latter, an elevated expression of genes involved in cholesterol metabolism was previously observed in oligodendroglial lineage cells during remyelination (Voskuhl et al., 2019). For instance, dietary cholesterol and 8,9-unsaturated sterols enhance the differentiation of OPCs, thereby promoting remyelination (Berghoff et al., 2017; Hubler et al., 2018). Furthermore, elevated cholesterol uptake promotes mTOR kinase activity in OPCs, thereby enhancing cholesterol synthesis and their maturation (Mathews and Appel, 2016). Finally, decreased cholesterol levels in the CNS of MS patients are associated with severe demyelination, which might well relate to perturbed remyelination in these patients (Evangelopoulos et al., 2022). How cholesterol mediates OPC maturation and remyelination remains poorly understood. However, there is now evidence that its mode of action relies on the lesion stage. While acute demyelinated lesions seem to rely on cholesterol recycling to enhance remyelination, chronic demyelinated lesions depend on increased local cholesterol synthesis in order to drive CNS repair (Berghoff et al., 2021a; Berghoff et al., 2021b). Given the abundance of macrophages in acute demyelinated lesions, this concept provides a molecular rationale for EVs acting as cholesterol traffickers and promoting endogenous OPC maturation and remyelination in active lesions, which often show signs of repair (Brück et al., 1997; Brück et al., 2003; Erickson, 2008; Staugaitis et al., 2012). It also argues for cholesterol-rich EVs, such as those released by RAMs, being of therapeutic interest to boost remyelination, especially in chronic lesions which are generally devoid of macrophages and microglia (Hagemeier et al., 2012; Hanafy and Sloane, 2011; Wolswijk, 2002).

Horizontal lipid flux is essential in supplying OPCs with cholesterol for developmental myelination and remyelination (Berghoff et al., 2022). With respect to the latter, horizontal transfer of recycled cholesterol from astrocytes to OLNs via lipoproteins is a major feature of normal brain development and myelination (Camargo et al., 2017; Werkman et al., 2021). Our findings now indicate that macrophage-derived EVs are vital traffickers of cholesterol in demyelinating disorders as well, thereby closely monitoring CNS repair. Accordingly, other studies defined the importance of EVs in trafficking of lipids in neurodegenerative disorders, with EV-associated monosialogangliosides, phospholipids, and ceramide analogues affecting amyloid-beta oligomerization, α-synuclein pathology, and neuroinflammation (Grey et al., 2015; Hoshino et al., 2013; Ikeda et al., 2011; Takasugi et al., 2015; Thomas and Salter, 2010). Guided by lipidomic analysis, microglia-derived EVs from Alzheimer’s disease patients were further shown to have increased levels of free cholesterol and decreased levels of docosahexaenoic acid-containing polyunsaturated fatty acids (Cohn et al., 2021). Finally, EV-associated sphingosine-1-phosphate derived from repair-associated microglia enhances OPC migration towards demyelinated lesions (Lombardi et al., 2019). While our findings identify EV-associated cholesterol as a key molecular determinant of the reparative features of RAMs, more research is warranted to assess whether other lipids, in concert with cholesterol, enhance OPC maturation and remyelination. Similarly, whereas our findings indicate that EVs released by RAMs directly impact OPC maturation *in vitro*, we cannot exclude that other cells, including astrocytes and neurons (Lombardi et al., 2019), control the regenerative impact of RAM-derived EVs on remyelination *ex vivo* and *in vivo*.

In summary, our findings indicate that EVs released by RAMs enhance OPC maturation and remyelination through EV-associated cholesterol, which is incorporated into the OPC membrane via direct membrane fusion. Our findings could help in designing novel therapeutic strategies, including the synthesis of cholesterol-enriched EVs or nanostructured lipid carriers, to support remyelination in demyelinating disorders.

## Acknowledgements

We thank L Timmermans, MP Tulleners, K Wauterickx, and M Jans for excellent technical assistance. The work was financially supported by the Research Foundation of Flanders (FWO Vlaanderen; 1S15519N, G099618FWO, and 12J9119N) and Interreg V-A EMR program (EURLIPIDS, EMR23). The funding agencies have no role in the design, analysis, and writing of the article.

## Author’s contributions

S. Vanherle, J.J.A. Hendriks, and J.F.J. Bogie conceived experiments. S. Vanherle, J. Guns, M. Loix, F. Mingneau, T. Dierckx, T. Vangansewinkel, P.P. Lins, J. Dehairs, and M. Haidar performed experiments. S. Vanherle, J. Guns, M. Loix, F. Mingneau, T. Dierckx, J. Dehairs, M. Haidar, and J.F.J. Bogie analyzed data. S. Vanherle, J. Guns, M. Loix, F. Mingneau, T. Dierckx, J.V.Swinnen, S.G.S. Verberk, M. Haidar, J.J.A. Hendriks, and J.F.J. Bogie discussed results. T. Vangansewinkel, E. Wolfs, P.P. Lins, A. Bronckaers, I. Lambrichts, J. Dehairs, and J.V. Swinnen contributed reagents, materials, and analysis tools. S. Vanherle, S.G.S. Verberk, J.J.A. Hendriks, and J.F.J. Bogie wrote the manuscript. S. Vanherle, J. Guns, M. Loix, F. Mingneau, T. Dierckx, T. Vangansewinkel, E. Wolfs, P.P. Lins, A. Bronckaers, I. Lambrichts, J. Dehairs, J.V. Swinnen, S.G.S. Verberk, M. Haidar, J.J.A. Hendriks, and J.F.J. Bogie revised the manuscript.

