## Supplemental figures for "Extracellular vesicle-associated cholesterol dictates the regenerative functions of macrophages in the brain"

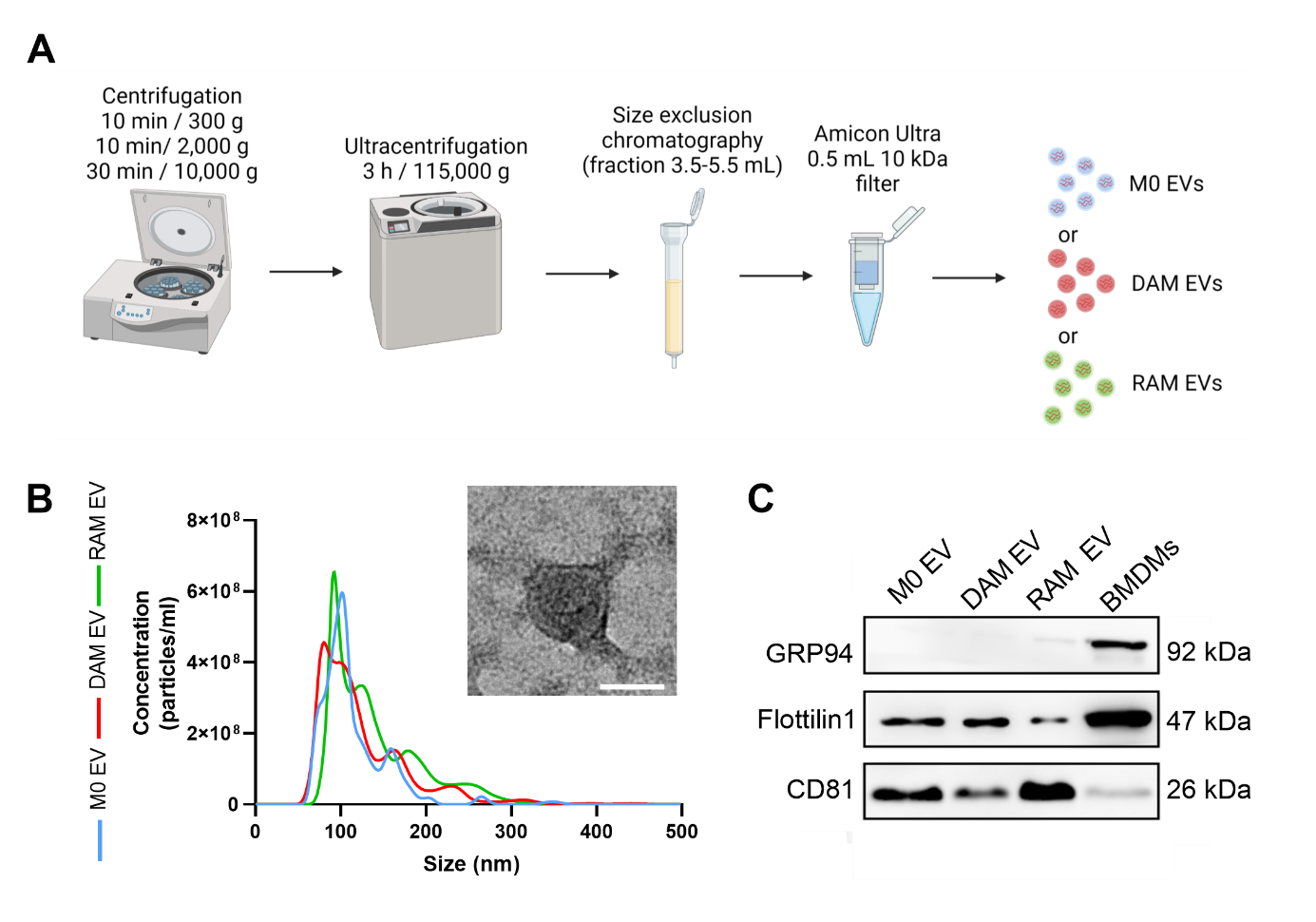

**Supplementary figure 1: Characterization of extracellular vesicles isolated from macrophages**. **A**) Schematic representation showing the isolation of extracellular vesicles (EVs) released by naive macrophages (M0, PBS-treated), disease-associated macrophages (DAMs, LPS-stimulated), and repair-associated macrophages (RAMs, IL-4-stimulated). Created with biorender.com. **B**) Particle size distribution and particles per mL of EVs isolated from M0, DAMs, and RAMs measured by Nanoparticle Tracking Analysis. Inset: representative transmission electron microscopy image of EVs isolated from macrophages. Scale bar, 50 nm. **C**) Immunoblot analysis of EV markers CD81 and Flottilin1, and GRP94 as negative EV marker of EVs isolated from M0, DAMs, and RAMs, and bone marrow-derived macrophage lysates.

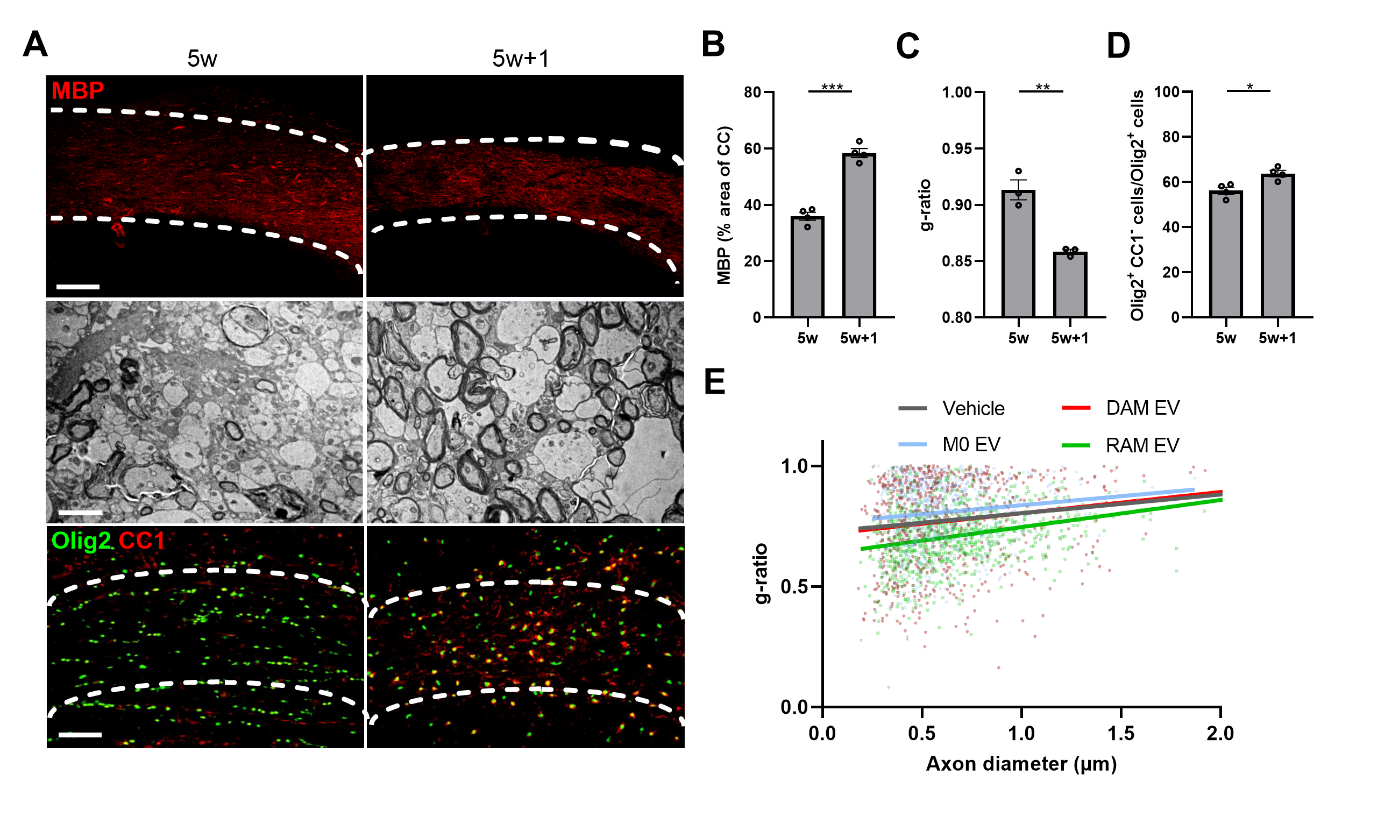

**Supplementary figure 2: Extracellular vesicles released by RAMs enhance remyelination in the cuprizone model. A**) Representative images of immunofluorescent MBP and Olig2/CC1 stains and transmission electron microscopy analysis of the corpus callosum (CC) from mice after demyelination (5w) and during remyelination (5w+1). The outer border of the CC is demarcated by the dotted line. Scale bar, 200 µm (rows 1, 3) and 2 µm (row 2). **B**) Quantification of the MBP^+^ area of the CC from cuprizone mice after demyelination (5w) and during remyelination (5w+1) (n = 3 animals, 3 images/animal). **C**) Analysis of the g-ratio (the ratio of the inner axonal diameter to the total outer diameter in CC from cuprizone mice after demyelination (5w) and during remyelination (5w+1) (n = 3 animals, 3 images/animal, 100-150 axons/image). **D**) Quantification of the percentage Olig2^+^ CC1^+^ cells out of total Olig2^+^ cells in the CC of cuprizone animals after demyelination (5w) and during remyelination (5w+1) (n = 3 animals, 3 images/animal). **E**) Analysis of g-ratio as a function of axon diameter in CC from cuprizone mice intracerebroventricularly treated with vehicle (PBS) or extracellular vesicles (EVs) released by naive macrophages (M0, PBS-treated), disease-associated macrophages (DAMs, LPS-stimulated), and repair-associated macrophages (RAMs, IL-4-stimulated). Mice were injected with vehicle or EVs (1 x 10^9^ EVs) after demyelination (5w), and analysis was done during remyelination (5w+1) (n = 3-5 animals, 3 images/animal, 100-150 axons/image). Data are represented as mean $\pm$ SEM and statistically analyzed using the Mann-Whitney test. *, p < 0.05; **, p < 0.01; ***, p < 0.001.

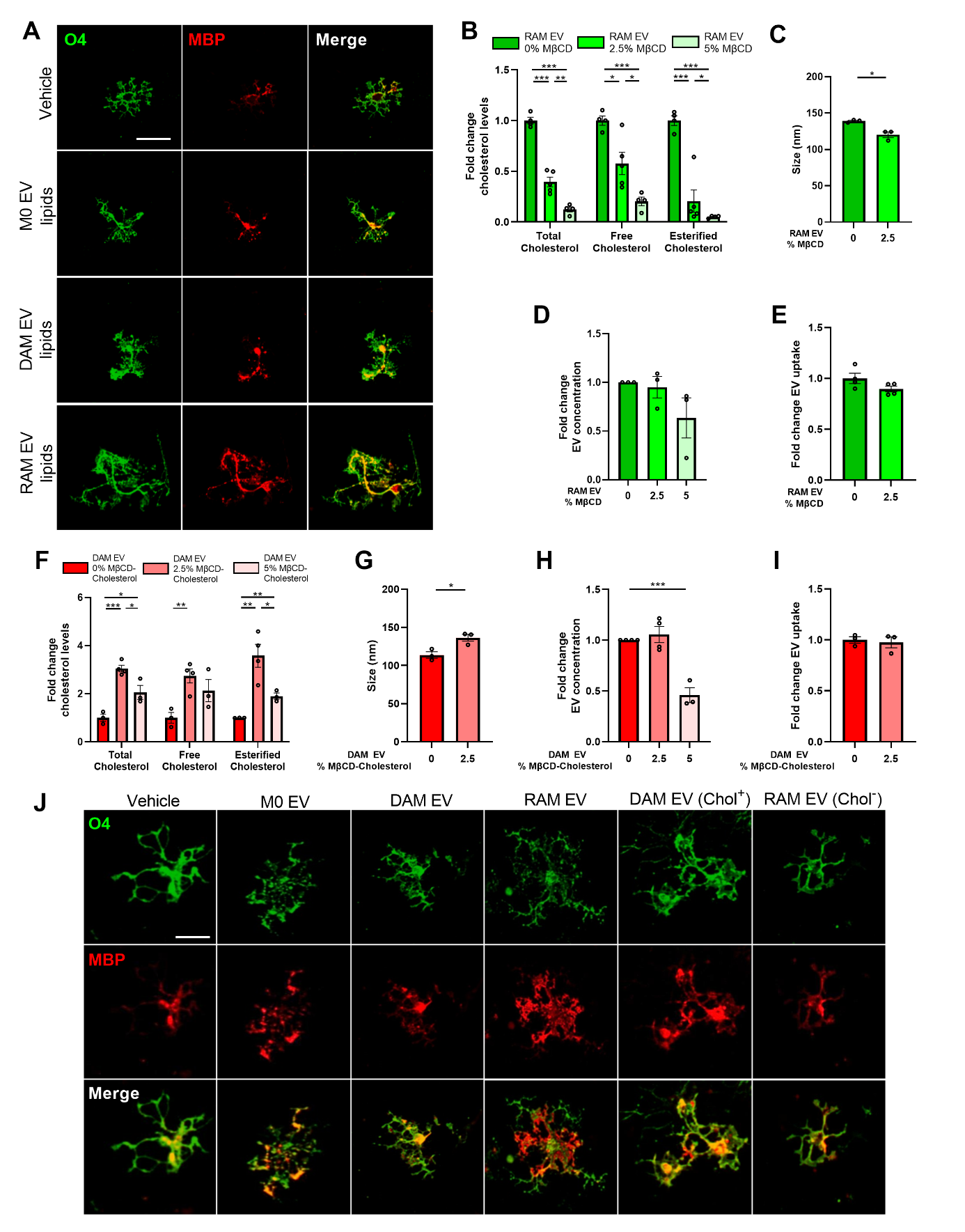

**Supplementary figure 3: Cholesterol abundance controls the impact of EVs released by macrophages on OPC maturation.** **A**) Representative images of immunofluorescent MBP (mature oligodendrocyte) and O4 (premature oligodendrocyte) stains of OPCs treated with lipids isolated from extracellular vesicles (EVs) released by naive macrophages (M0, PBS-treated), disease-associated macrophages (DAMs, LPS-stimulated), and repair-associated macrophages (RAMs, IL-4-stimulated). Scale bar, 25 µm. **B,F**) Quantification of total, free, and esterified cholesterol in EVs released by RAMs and DAMs (n = 3-4 isolates), which were depleted of or enriched with cholesterol, respectively. Cells were exposed to cholesterol-loaded or unloaded methyl-beta cyclodextrin (MβCD; 0% m/v, 2.5% m/v, or 5% m/v). **C, D,** **G, H**) Nanoparticle Tracking Analysis to assess the size (C,G) and concentration (D,H) of EVs released by of RAMs and DAMs that were exposed to 0%, 2.5% , or 5% MβCD (n = 3 isolates). **E,I**) Internalization of DiI-labeled EVs released by untreated and cholesterol-enriched DAMs (E), as well as untreated and cholesterol-depleted RAMs (I), by OPC. OPCs were exposed for 3 h to 4 x 10^8^ EVs/mL. **J**) Representative images of immunofluorescent MBP (mature oligodendrocyte) and O4 (premature oligodendrocyte) stains of OPCs treated with EVs released by M0 macrophages, DAMs, and RAMs, as well as EVs released by DAMs and RAMs that were enriched (Chol^+^) with or depleted (Chol^-^) of cholesterol, respectively. OPCs were treated with 4 x 10^8^ EVs/mL. Scale bar, 25 µm. Results are pooled from three biological replicates. Data are represented as mean $\pm$ SEM and statistically analyzed using the Kruskal-Wallis test followed by Dunn’s multiple comparison test. *, p < 0.05; **, p < 0.01; ***, p < 0.001.

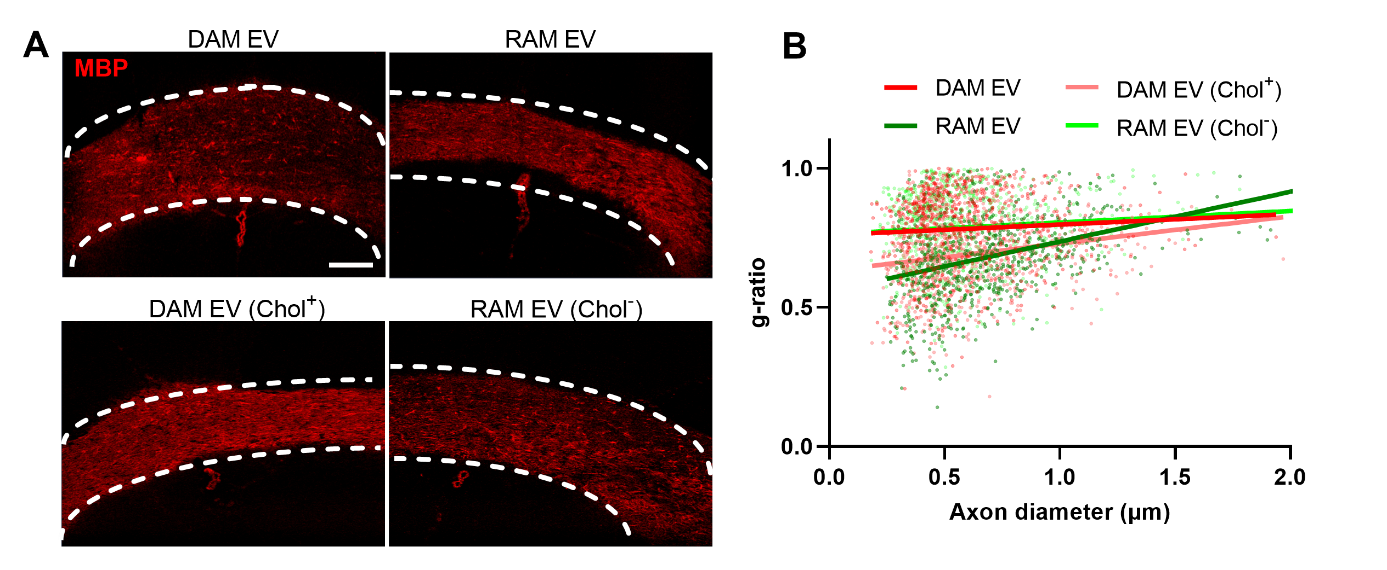

**Supplementary figure 4: Cholesterol abundance controls the impact of EVs released by macrophages on remyelination. A**) Representative images of immunofluorescent MBP stain of the corpus callosum (CC) from mice treated intracerebroventricularly with vehicle or EVs released by disease-associated macrophages (DAMs, LPS-stimulated), and repair-associated macrophages (RAMs, IL-4-stimulated), as well as EVs released by DAMs and RAMs that were enriched (Chol^+^) with or depleted (Chol^-^) of cholesterol, respectively. The outer border of the CC is demarcated by the dotted line. Mice were intracerebroventricularly injected with vehicle or EVs (1 x 10^9^ EVs) after demyelination (5w), and analysis was done during remyelination (5w+1). **B**) Analysis of g-ratio as a function of axon diameter in CC from cuprizone mice treated with vehicle or divergent EVs during remyelination (5w+1) (n = 5-6 animals, 3 images/animal, 100-150 axons/image).

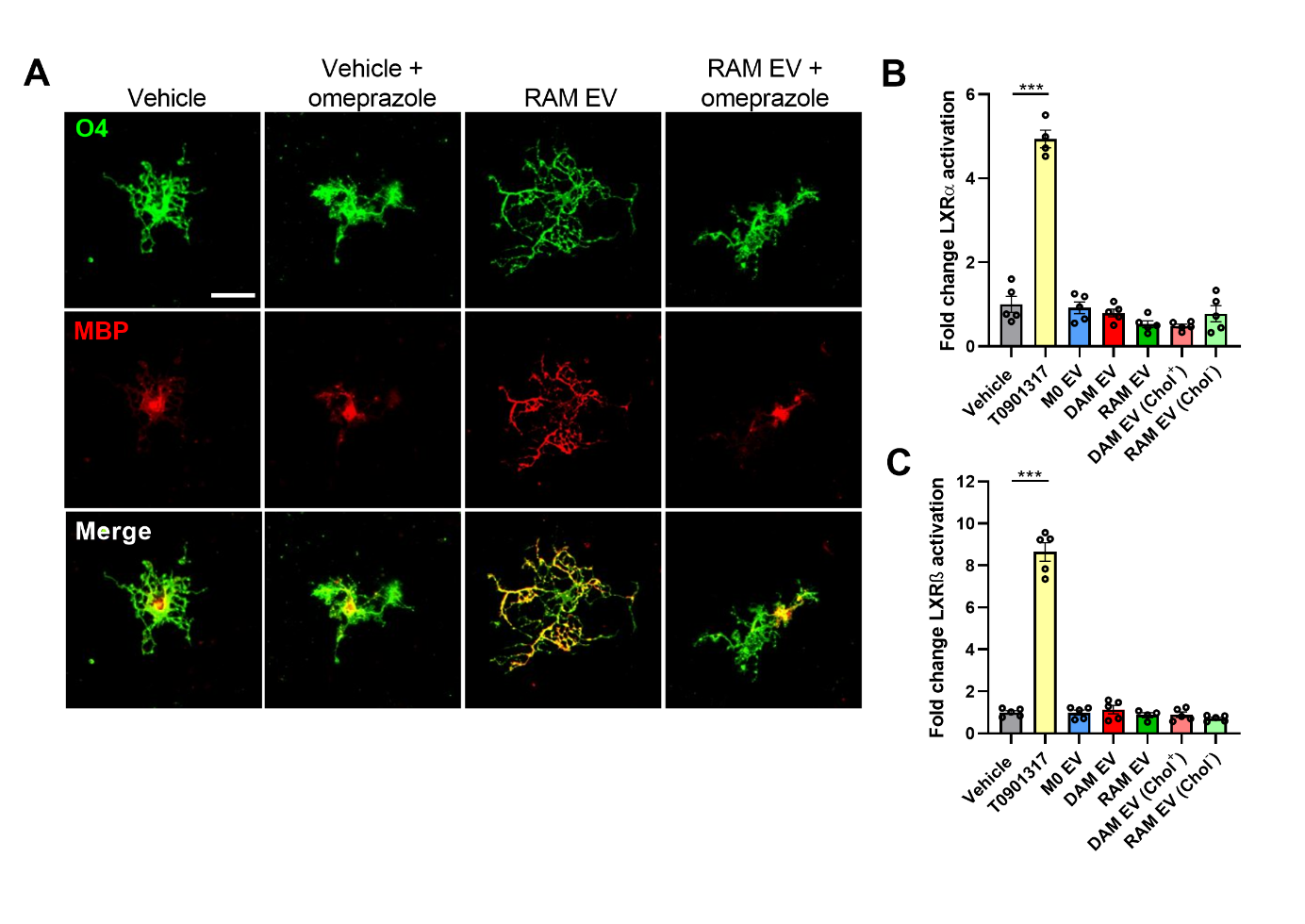

**Supplementary figure 5: Extracellular vesicle-associated cholesterol promotes OPC maturation via direct membrane fusion independent of LXR activation. A**) Representative images of immunofluorescent MBP (mature oligodendrocyte) and O4 (premature oligodendrocyte) stains of OPCs treated with vehicle or extracellular vesicles (EVs) released by repair-associated macrophages (RAMs, IL-4-stimulated), and/or omeprazole. OPCs were treated with 4 x 10^8^ EVs/mL and 50 µM omeprazole. Scale bar, 25 µm. **B-C**) Liver X receptor (LXR) α (B) and β (C) ligation in COS7 cells following 24 h exposure to the LXR agonist T0901317, EVs released by naive macrophages (M0, PBS-treated), disease-associated macrophages (DAMs, LPS-stimulated), and RAMs, as well as EVs isolated from DAMs and RAMs that were enriched (Chol^+^) with or depleted (Chol^-^) of cholesterol, respectively. LXRα and LXRβ activation was defined by a luciferase-based nuclear receptor reporter assay (n = 5 wells). OPCs were treated with 4 x 10^8^ EVs/mL and 10 µM T0901317. All results are pooled from three biological replicates. Data are represented as mean $\pm$ SEM and statistically analyzed using the Kruskal-Wallis test followed by Dunn’s multiple comparison test (B-C). ***, p < 0.001.
